# Activation of the integrative and conjugative element Tn*916* causes growth arrest and death of host bacteria

**DOI:** 10.1101/2022.04.01.486793

**Authors:** Emily L. Bean, Lisa K. McLellan, Alan D. Grossman

## Abstract

Integrative and conjugative elements (ICEs) serve as major drivers of bacterial evolution. These elements often confer some benefit to a host cell, including antibiotic resistance, metabolic capabilities, or pathogenic determinants. ICEs can also have negative impacts on their host cells. Here, we investigated the effects of the ICE (conjugative transposon) Tn*916* on host cells. Because Tn*916* is active in a relatively small subpopulation of host cells, we developed a fluorescent reporter system for monitoring activation of Tn*916* in single cells. We found that when active in *Bacillus subtilis* and its natural host *Enterococcus faecalis*, Tn*916* inhibited cell division and most cells died. We also observed these phenotypes on the population level in *B. subtilis* utilizing a modified version of Tn*916* that can be activated in the majority of cells. We identified two genes (*orf17* and *orf16*) in Tn*916* that were sufficient to cause host growth defects and identified a single gene, *yqaR*, that is found in a defective phage (*skin*) in the *B. subtilis* chromosome that is required for this phenotype. However, these three genes are only partially responsible for the growth defect caused by Tn*916*, indicating that Tn*916* possesses multiple mechanisms to affect growth and viability of host cells. These results highlight the complex relationships that conjugative elements have with their host cells and the interplay between mobile genetic elements.

## INTRODUCTION

Integrative and conjugative elements (ICEs), also called conjugative transposons, are a type of mobile genetic element that serve as drivers of bacterial evolution. Typically, they reside integrated in a bacterial host chromosome. Either stochastically, or in response to a signal, they can excise from the chromosome forming a plasmid. ICE-encoded conjugation machinery (a type IV secretion system, T4SS) can transfer the ICE into a recipient cell in a contact-dependent manner (Bellanger et al., 2014; Delavat et al., 2017; Johnson and Grossman, 2015; Roberts and Mullany, 2009; Wozniak and Waldor, 2010).

Conjugative elements often carry genes that confer phenotypes to host cells, including antibiotic resistances, pathogenic or symbiotic abilities, and various metabolic capabilities. Conjugative elements were initially identified based on the phenotypes that they confer to bacterial hosts (Frost et al., 2005; Johnson and Grossman, 2015; Treangen and Rocha, 2011). Advantageous phenotypes conferred by ICEs likely mitigate potential costs of maintaining these elements.

Conjugative elements can also have more complex relationships with their host cells. Some elements encode functions that manipulate host development, growth, and viability (for examples see (Beaber et al., 2004; Jones et al., 2021; Pembroke and Stevens, 1983; Reinhard et al., 2013)). Excessive mating events can be detrimental to host viability (Avello et al., 2019; Skurray and Reeves, 1973). Additionally, interactions between conjugative elements and other horizontally-acquired elements, like phages, can impact a host cell. For instance, T4SSs can be targeted by male-specific phages (Caro and Schnös, 1966; Lang et al., 2014; Loeb, 1960), phages can prevent conjugation events from occurring (Lin et al., 2011; Novotny et al., 1968; Ou, 1973), or ICEs can encode abortive infection mechanisms to prevent phage production (Johnson et al., 2022).

Here, we present evidence that the ICE Tn*916* possesses a previously unknown ability to cause a growth arrest and kill its host cell. Tn*916* was the first ICE discovered and was identified based on its ability to spread tetracycline resistance between two strains of *Enterococcus faecalis* (Franke and Clewell, 1981a, 1981b). Tn*916* and its relatives have since been found in other Gram-positive bacteria including *Streptococcus, Staphylococcus*, and *Clostridium* species (Clewell and Flannagan, 1993; Clewell et al., 1985; Fitzgerald and Clewell, 1985; Roberts and Mullany, 2009, 2011; Sansevere and Robinson, 2017; Santoro et al., 2014), and it is functional in *Bacillus subtilis* (Christie et al., 1987; Ivins et al., 1988; Mullany et al., 1990; Roberts et al., 2003; Scott et al., 1988; Wright and Grossman, 2016). Tn*916* is regulated, at least in part, by a transcriptional attenuation mechanism that is relieved in the presence of tetracycline or other translation-inhibiting drugs (Celli and Trieu-Cuot, 1998; Roberts and Mullany, 2009; Scornec et al., 2017; Su et al., 1992). These drugs stimulate excision and transfer of Tn*916* (Celli and Trieu-Cuot, 1998; Manganelli et al., 1995; Showsh and Andrews, 1992; Wright and Grossman, 2016). However, Tn*916* only activates and excises in ∼0.1-3% of a population of host cells (Celli and Trieu-Cuot, 1998; Celli et al., 1997; Marra and Scott, 1999; Scott et al., 1994; Wright and Grossman, 2016). Therefore, any effects Tn*916* activation has on the host cell would be masked in population level analyses.

To study the effects of Tn*916* gene activation on the population-level in *B. subtilis* host cells, we used a hybrid conjugative element that contains the regulatory and recombination genes from a heterologous element and the DNA processing and conjugation genes from Tn*916*. Using this hybrid element, we identified two Tn*916* genes that are sufficient to cause *B. subtilis* host cells to stop growing. We also identified a gene in the defective phage *skin* that was required for the growth defects caused by the two Tn*916* genes.

We also analyzed the effects of Tn*916* on cell growth in single cells using a fluorescent reporter to monitor activation of Tn*916*. We found that cell growth and division was inhibited in cells with an activated (excised) Tn*916*. Furthermore, most of these cells died. When activated in its natural host, *E. faecalis*, Tn*916* also caused growth arrest and cell death. We suggest that these growth defects may be a common feature across other bacterial hosts of Tn*916* and Tn*916*- like elements. Our results also indicate that the growth arrest likely functions to limit the spread of the element.

## RESULTS

### Increased activation of Tn*916* genes causes defects in cell growth and viability

Tn*916*, like many conjugative elements, only becomes active and excises from the genome in a small portion (∼0.1-3%) of the cells in a population (Celli and Trieu-Cuot, 1998; Celli et al., 1997; Marra and Scott, 1999; Scott et al., 1994; Wright and Grossman, 2016). In previous work, we created hybrid ICEs that contained the DNA processing and conjugation functions of Tn*916* and the efficient regulatory and recombination (integration and excision) systems encoded by ICE*Bs1* (Bean et al., 2021). We refer to this hybrid element as (ICE*Bs1*-Tn*916*)-H1, or H1 for short (Figure 1). ICE*Bs1* and H1 can be activated in ∼25-90% of the cells in a population by overproduction of its activator protein RapI (Auchtung et al., 2005, 2007; Bean et al., 2021; Lee et al., 2007). The ability to activate Tn*916* genes needed for DNA processing and conjugation in a large proportion of cells enables population-level analyses of effects of these genes on host cells.

**Figure 1.**
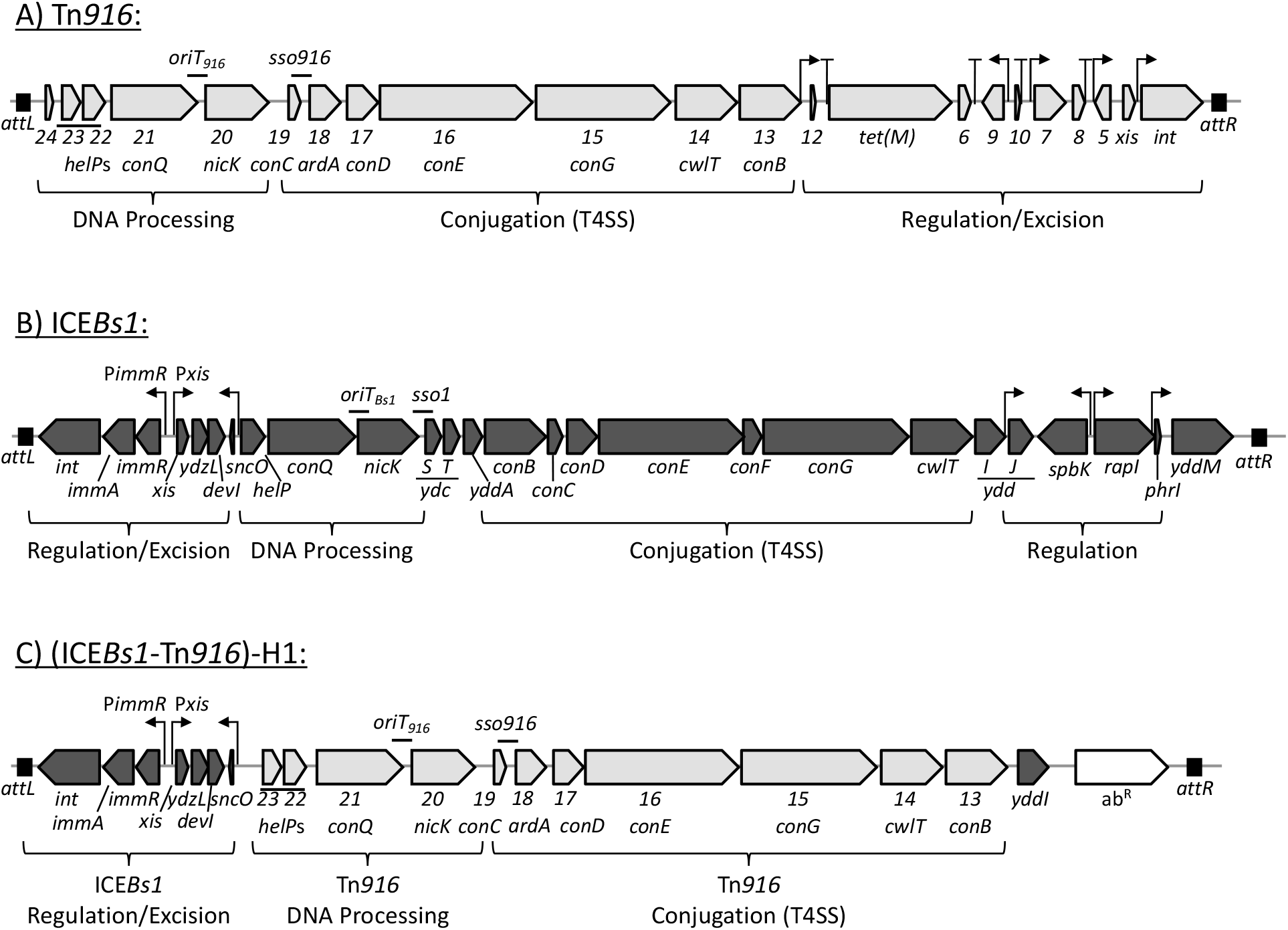
Genetic maps of Tn*916*, ICE*Bs1*, and (ICE*Bs1*-Tn*916*)-H1. Maps of the conjugative elements used in these studies are shown: **A)** Tn*916*, **B)** ICE*Bs1*, and **C)** (ICE*Bs1*- Tn*916*)-H1. Open reading frames are indicated by horizontal arrows (gray for Tn*916*, black for ICE*Bs1*). Tn*916* gene names are abbreviated to include only the number designation from the name (ie “*orf23”* is written as “*23*”), and the corresponding ICE*Bs1* homolog gene name is written in parentheses below, when appropriate. H1 contains a combination of Tn*916* and ICE*Bs1* genes, as previously reported (Bean et al., 2021). Functional modules are indicated by brackets below each map. Black boxes indicate attachment sites *attL* and *attR*: Tn*916* is integrated between *yufK* and *yufL*, unless otherwise indicated; ICE*Bs1* and H1 are integrated at *trnS*-*leu2*. Promoters are indicated by bent arrows; putative transcription terminators are indicated by “T” shapes. The current model of transcriptional regulation of Tn*916* shown in (b) is adapted from (Roberts and Mullany, 2009; Su et al., 1992). Previously determined origins of transfer (*oriT*) and single strand origins of replication (*sso*) are indicated by a “-“ above the genetic map (Jaworski and Clewell, 1995; Lee and Grossman, 2007; Wright and Grossman, 2016; Wright et al., 2015).

We found that there was a dramatic drop in viability of cells containing an activated H1 compared to that of cells containing an activated ICE*Bs1* or Tn*916*. We monitored the growth and viability of host cells that were grown in defined minimal medium under activating and non-activating conditions: xylose was added to induce expression of *rapI* and activation of either H1 (ELC1214) or ICE*Bs1* (MMB970); tetracycline (2.5 µg/ml) was added to stimulate Tn*916* activation. By two hours after induction, excision had occurred in ∼70%, ∼95%, and 0.1% of cells containing H1, ICE*Bs1*, and Tn*916*, respectively.

Approximately one hour after induction of H1, growth of the culture stopped as measured by optical density (Figure 2A). The optical density of the culture then declined (Figure 2B), indicating that cell lysis was likely occurring. Indeed, there was a drop in viable cells in the culture in which H1 was activated. Cultures in which H1 was induced had an ∼100-fold drop in viable cells relative to the uninduced culture (Figure 2B), as measured by colony forming units (CFUs) three hours after activation of the element. Furthermore, the final number of viable cells after activation was ∼2% of that before activation of H1. Together, these results indicate that activation of H1 caused a growth arrest and cell death.

**Figure 2.**
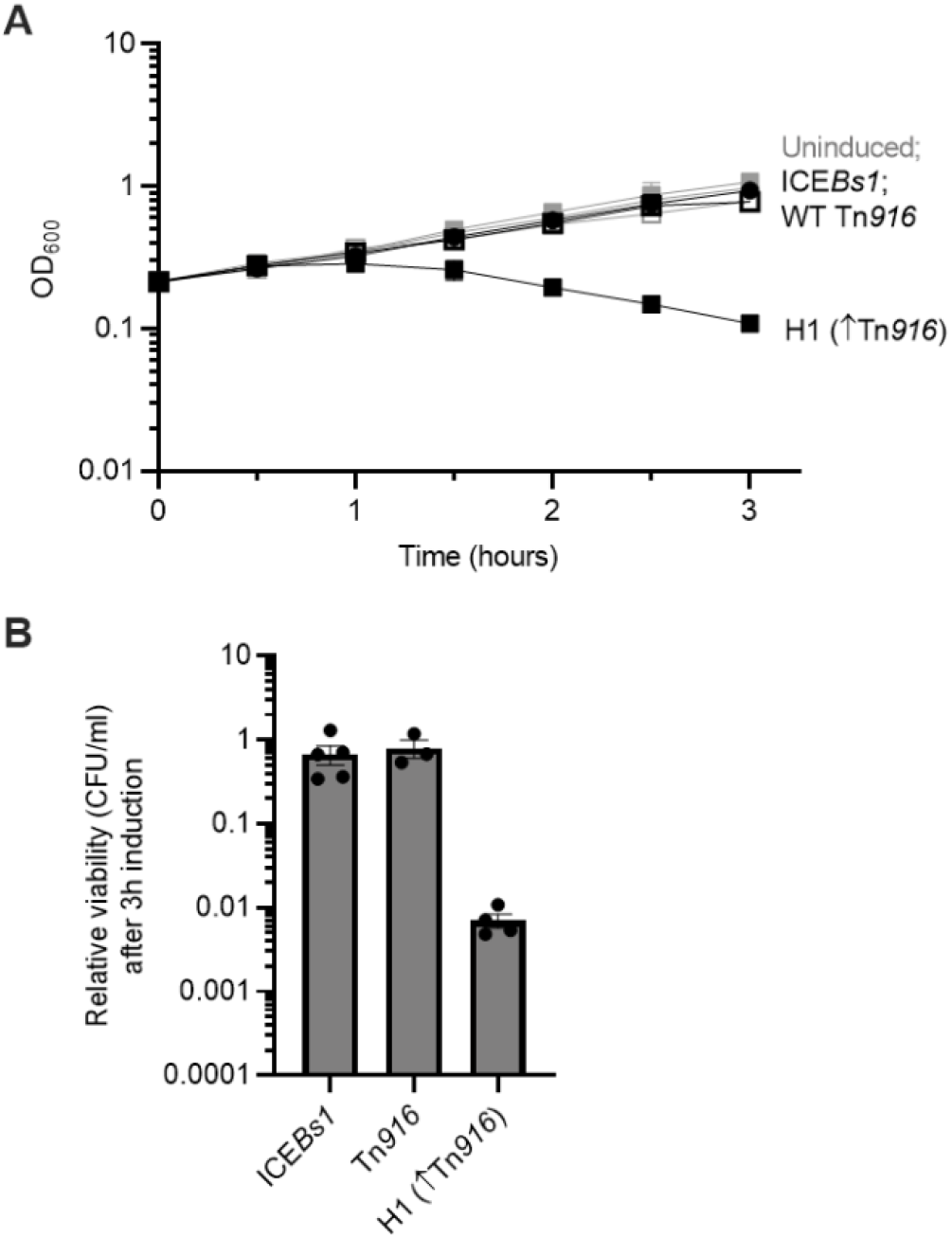
H1 (↑Tn*916*) activation causes a growth arrest and cell death. Strains containing ICE*Bs1* (circles, MMB970), Tn*916* (open squares, CMJ253) or H1 (closed squares, ELC1214) were grown in minimal arabinose medium to early exponential phase. At time = 0 hours, when cultures were at an OD_600_ ∼0.2, cultures were split into inducing (+1% xylose to stimulate *rapI* expression, or +2.5 µg/ml tetracycline to stimulate Tn*916* activation) and non- inducing conditions. **A)** Growth was monitored by OD_600_ for three hours. Black lines indicate growth of the indicated induced cultures; gray lines indicate growth of uninduced cultures. **B)** The relative viability of cultures after three hours of element induction was calculated as the number of colony forming units (CFUs) formed by the induced culture, divided by that from the uninduced culture (a value of “1” indicates there is no change in CFUs with induction). Data presented are averages from three or more independent experiments (individual data points shown in (B)), with error bars depicting standard error of the mean. Error bars could not always be depicted in (A) due to the size of each data point.

In contrast, cells in which ICE*Bs1* had been induced continued to grow, plateaued at a relatively normal optical density (Figure 2A), and there was no evidence of a large drop in cell viability (Figure 2B). Cultures of Tn*916*-containing cells also grew normally (Figure 2A) and there was no apparent drop in cell viability (Figure 2B). Of course, even if all of the cells (∼0.1%) in which Tn*916* had become activated had lost viability, we would not detect this on a population level with the assays used (Figure 2). Based on the effects of H1 and ICE*Bs1* on cell growth and viability, we infer that the defects caused by H1 were either due to increased expression of genes from Tn*916* or the absence of a protective gene(s) from ICE*Bs1* in the ICE*Bs1*-Tn*916* hybrid.

We found that the growth defects caused by induction of H1 were not due to loss of some putative protective gene(s) in ICE*Bs1*. We used a mutant of ICE*Bs1* that contained only the ICE*Bs1* genes present in H1. That is, the mutant {ICE*Bs1* (Δ*helP*-Δ*cwlT*, Δ*yddJ*-*yddM*); strain ELC1226} was missing all the ICE*Bs1* genes that were also missing in H1, and also did not contain any genes from Tn*916*. Activation of this element did not cause a growth defect (Figure S1), indicating that the growth defect was not due to the absence of some protective ICE*Bs1* gene(s).

### *orf16* and *orf17* in Tn*916* are partially required for the growth arrest caused by activation of the ICE*Bs1*-Tn*916* hybrid H1

We made deletions in H1 to determine which of the Tn*916* genes were responsible for the growth defect. Initially, this analysis was complicated by loss of H1 in a proportion of host cells after activation. Approximately 32% of H1 host cells post-induction did not contain the element, as determined by checking 100 colonies for kanamycin resistance (which is encoded on H1). For this reason, we used an excision-defective derivative of H1 in subsequent experiments. The excision-defective versions lack *attR*, the “right” attachment site that is necessary for element excision. The genes encoding the relaxases (*nicK* from ICE*Bs1*, *orf20* from Tn*916* in H1) were also removed to prevent the relaxase from nicking the host chromosome at the origin of transfer (*oriT*), which we have previously shown to negatively impact host cell viability (Lee et al., 2007; Menard and Grossman, 2013). For simplicity, we will refer to this element as H1-Δ*attR* (Materials and Methods).

Host strains containing excision-defective copies of H1 with single and pairwise deletions of Tn*916* genes were used to identify *orf16* and *orf17* as contributors to the growth arrest and killing phenotypes. Under identical conditions to those described above, we monitored the growth of host strains containing a xylose-inducible copy of H1-Δ*attR* (ELC1076), or H1-Δ*attR* with in the following deletions: *Δorf17* (ELC1419), Δ*orf16* (ELC1420), Δ*orf17*-*16* (ELC1942) under element activating and non-activating conditions. In parallel, a host strain containing an excision-defective copy of ICE*Bs1* (ELC1095) was used as a control strain that should not undergo a growth arrest following element activation.

### The excision-defective copy of H1 elicited a greater decrease in host viability than its excision-competent parent element

The H1-Δ*attR* host strain began exhibiting a decrease in growth as determined by OD approximately one hour post-induction, a similar timescale to the parent element (Figure 3A). However, three hours after induction, the relative viability of cells with H1-*ΔattR* was ∼2000- fold lower than that of cells with the uninduced element (Figure 3B). In contrast, the viability of cells with H1 (capable of excision) was ∼100-fold lower than that of that of cells with the uninduced element (Figure 2B) The ∼20-fold increase in the viability of cells in which the element can excise is likely due, at least in part, to the loss of the plasmid form of the element and then the selective advantage of cells without an active element.

**Figure 3.**
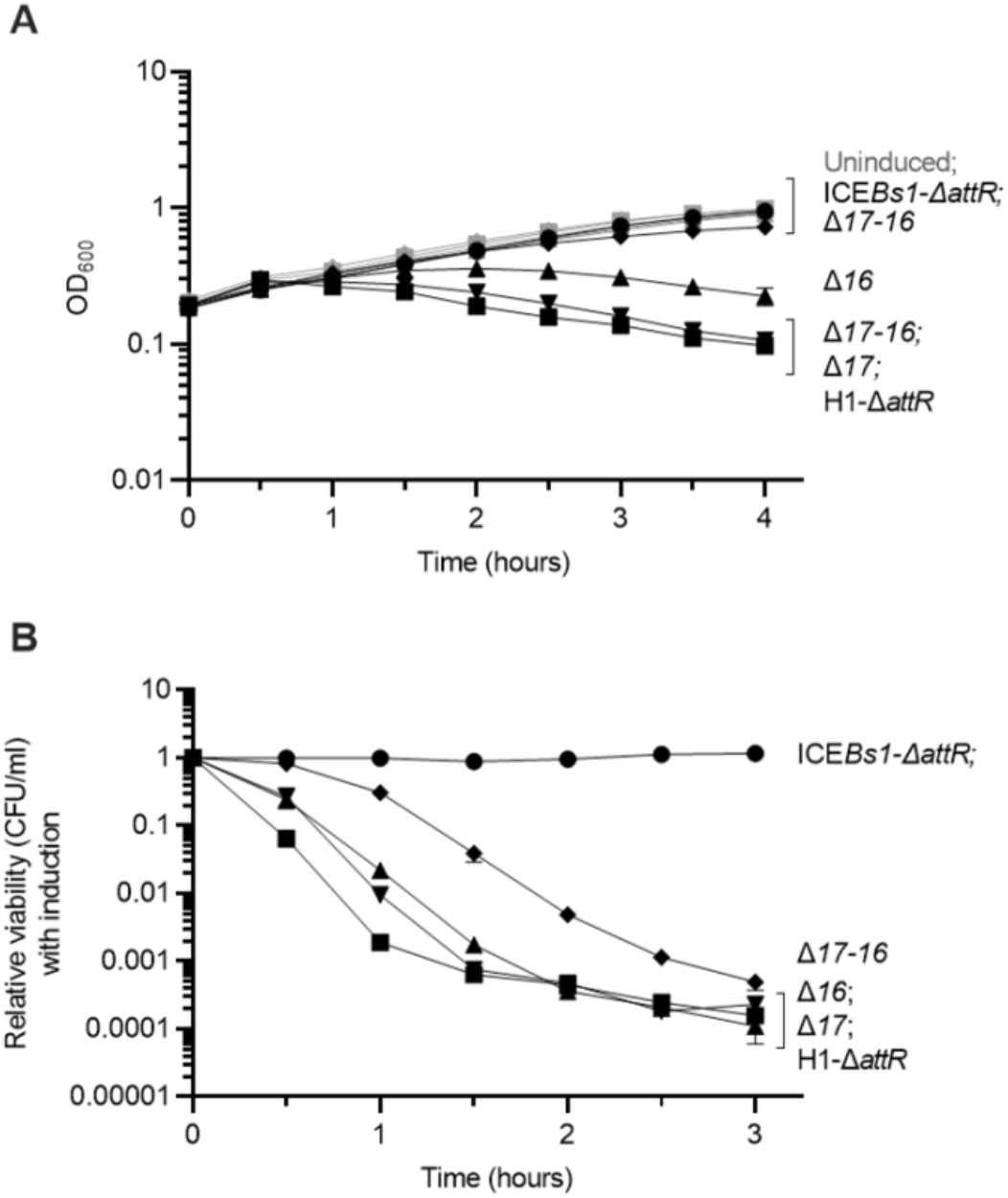
*orf16* and *orf17* are involved in the Tn*916*-caused growth arrest. Strains containing ICE*Bs1*-Δ*attR* (circles, ELC1095), or H1-Δ*attR* (squares, ELC1076) with the indicated deletion: Δ*orf17* (downward triangle, ELC1419), Δ*orf16* (upward triangle, ELC1420), and Δ*orf17*-*16* (diamonds, ELC1942) were grown in minimal arabinose medium to early exponential phase. At time = 0 hours, when cultures were at an OD_600_ ∼0.2, cultures were split into inducing (+1% xylose to stimulate *rapI* expression) and non-inducing conditions. **A)** Growth was monitored by OD_600_ for three hours. Black lines indicate growth of the indicated induced cultures; gray lines indicate growth of uninduced cultures. **B)** The relative viability of cultures was evaluated every 30 minutes for three hours post- induction and was calculated as the number of colony forming units (CFUs) formed by the induced culture, divided by that from the uninduced culture (a value of “1” indicates there is no change in CFUs with induction). Data presented are averages from three or more independent experiments (individual data points shown in (B)), with error bars depicting standard error of the mean. Error bars could not always be depicted in (A) and (C) due to the size of each data point.

We found that deletion of *orf16* (*virB4*-like; homolog of *conE* in ICE*Bs1*) and/or *orf17* (*virB3*-like; homolog of *conD* in ICE*Bs1*) partly suppressed the growth defect and drop in viability due to the activation of H1-Δ*attR* (Figure 3A, 3B). Deleting *orf16* improved the ODs of cultures at all time points post-induction, although a plateau was reached at approximately two hours, followed by a steady decline. There was a greater improvement when both *orf16* and *orf17* were deleted simultaneously: the ODs of these cultures continued increasing and were nearly indistinguishable from the uninduced cultures until three hours post-induction, at which point the uninduced cultures were slightly higher. However, deleting *orf17* alone only modestly improved the ODs of cultures. At three hours post-induction, the ODs of Δ*orf17* cultures were indistinguishable from H1-Δ*attR*.

Deletion of both *orf16* and *orf17* also partially rescued the viability defect as measured by CFUs. We examined relative CFU recovery every 30 minutes for three hours post-induction. Δ*orf17*-*16* improved recovered CFUs at all time points tested relative to H1-Δ*attR*, by up to ∼100-fold at one hour post-induction. This disparity decreased as the induction period went on. On the contrary, cells with deletion of either *orf16* or *orf17* alone had similar viability (CFUs) to cells with H1-Δ*attR* (Figure 3B). Δ*orf16* and Δ*orf17* each led to a ∼5-10 fold improvement in recovered CFUs compared to H1-Δ*attR* at 0.5 and one hour post-induction (Figure 3B). After this point, the number of recovered CFUs was nearly indistinguishable among these strains. Of note, we do not believe the improvement in Δ*orf17*-*16* viability is due to polar effects of the deletion on downstream genes; when Δ*orf17*-*16* was complemented with an inducible copy of *orf17*-*16* at an ectopic site (*lacA*::P*xis orf17*-*orf16*, ELC1938), the growth defects were completely restored to the levels exhibited by H1-Δ*attR* (Figure S1).

We hypothesized that any effect Orf16 is having on the host cell could be due to its activity as a VirB4-like ATPase. We monitored the impact of activation of an element containing a mutation in the predicted Walker A motif of Orf16 (K477E). Although this mutation abolishes conjugative transfer of Tn*916* and H1, the phenotype caused by the point mutation with respect to cell growth was indistinguishable from that of wild type *orf16* (Figure S1).

We also monitored the effects of all other Tn*916* gene deletions on host growth. No other deletion improved the optical density of the culture following element activation, but strains deleting the coupling protein (*orf21; virD4*-like; *conQ* in ICE*Bs1*), two of the essential components of the conjugation machinery (*orf19* (no *vir* counterpart; same predicted topology as *conC* in ICE*Bs1*), or *orf15* (*virB6*-*like*; *conG* in ICE*Bs1*)) all recovered ∼10-fold more CFUs three hours post-induction than the parent strain (Figure S1). Critically, no single gene deletion fully restored growth of cells containing this element, indicating that multiple Tn*916* genes are responsible for these growth defects. Altogether, these results indicated that Orf16, and to some extent Orf17, are partially responsible for Tn*916*-caused growth defects upon element activation. However, multiple other Tn*916* genes must be responsible for the remaining growth defect phenotypes. We hypothesize that Tn*916* encodes multiple mechanisms to negatively impact host cell growth and viability and decided to focus on the effects of Orf16 and Orf17 for the rest of this work.

### *orf17* and *orf16* together are sufficient to cause a growth arrest

We found that induction of *orf17* and *orf16* together was sufficient to cause a growth arrest in the absence of other Tn*916* ICE genes. In strains devoid of any ICEs, we placed *orf17*, *orf16*, or *orf17*-*16* together under the regulatory control of P*xis* from ICE*Bs1* at an ectopic site, *lacA* (strains ELC1494, ELC1491, and ELC1496, respectively). An empty expression construct (ELC1495) was used as a control. These strains all contained ectopic copies of the genes required for regulation of P*xis* (*immR, immA*, and a xylose-inducible copy of *rapI*). To monitor the effects of activation of these genes, we used both OD measurements and relative CFU formation to assess growth and viability of the cultures (using identical growth conditions described above).

Inducing expression of *orf17* or *orf16* alone was not sufficient to impact growth more than the empty expression construct. The growth curves of these strains were indistinguishable under inducing vs. non-inducing conditions (Figure 4A). There was a ∼2-3 fold decrease in CFUs that were recovered from the induced *orf17*, *orf16*, and empty control cultures relative to the non-induced cultures after a three-hour induction indicating that there was some cost associated with activating this induction system. Together, these results indicated that inducing *orf17* or *orf16* individually does not impact cell growth.

**Figure 4.**
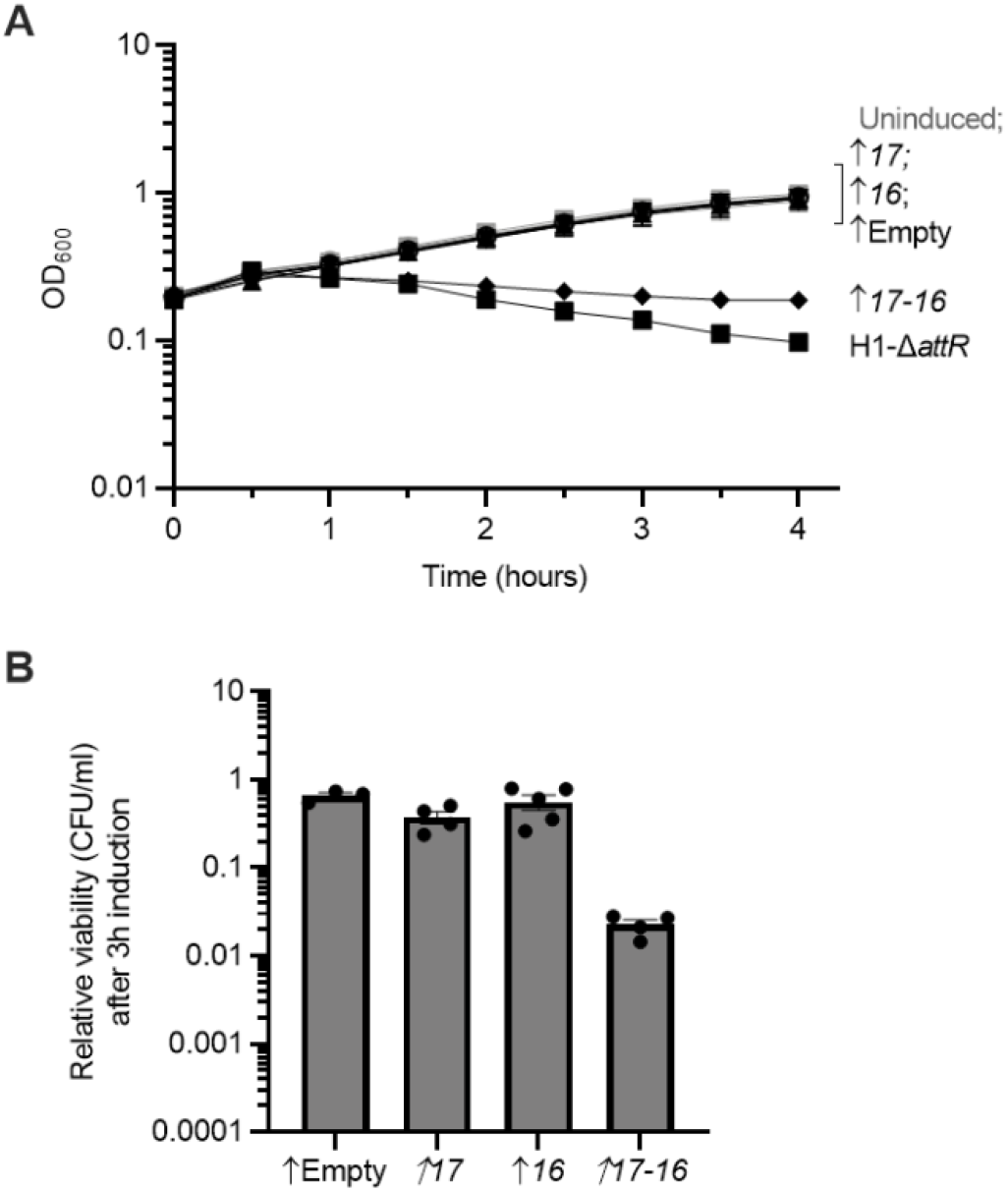
Orf17-16 are sufficient to cause a host cell growth arrest. Strains containing overexpression alleles of *orf17* (downward triangle, ELC1494), *orf16* (upward triangle, ELC1491), *orf17*-*16* (diamonds, ELC1496), or an empty cassette (open circles, ELC1495) and a strain containing H1-Δ*attR* (squares, ELC1076) were grown in minimal arabinose medium to early exponential phase. At time = 0 hours, when cultures were at an OD_600_ ∼0.2, cultures were split into inducing (+1% xylose to stimulate *rapI* expression) and non-inducing conditions. **A)** Growth was monitored by OD_600_ for three hours. Black lines indicate growth of the indicated induced cultures; gray lines indicate growth of uninduced cultures. Growth curves of ELC1076 (containing H1-Δ*attR*) from Figure 3A were included as reference. **B)** The relative viability of cultures after three hours of element induction was calculated as the number of colony forming units (CFUs) formed by the induced culture, divided by that from the uninduced culture (a value of “1” indicates there is no change in CFUs with induction). Data presented are averages from three or more independent experiments (individual data points shown in (B)), with error bars depicting standard error of the mean. Error bars could not always be depicted in (A) due to the size of each data point.

However, inducing expression of *orf17* and *orf16* together led to a decrease in OD after one hour (Figure 4A). This trend was similar to that observed with induction of all Tn*916* genes in the context of H1 (Figures 2A, 3A), however the drop in OD was less severe when only *orf17*-*16* were overexpressed. This difference further indicates that other Tn*916* genes contribute to the full effect on cell growth. Induction of *orf17* and *orf16* together also caused an approximately 50-fold drop in CFUs three hours after induction of expression (Figure 4B). These data indicate that Orf17 and Orf16 contribute to the drop in viability following induction of Tn*916* genes in H1, but that other Tn*916* genes are also required for the nearly 2000-fold drop in CFUs observed upon induction of Tn*916* genes in H1-Δ*attR*.

Together, these results indicate that Orf17 and Orf16, in the absence of any other Tn*916* gene products, are sufficient to cause a growth arrest and cell death of *B. subtilis*. We suspect that *orf17* is needed for the proper expression of *orf16*. This is by analogy to the homologous genes *conD* (*orf17*) and *conE* (*orf16*) in ICE*Bs1* where ectopic expression of *conE* (*orf16*) is improved in the presence of the upstream gene *conD* (*orf17*), likely due to translational coupling (Berkmen et al., 2010). Alternatively, both proteins may be required to interact with some host component. In the context of ICE*Bs1*, it has been shown that ConD assists in localizing ConE, a cytoplasmic protein, to the membrane (Leonetti et al., 2015). Perhaps a similar interaction is occurring in the context of Tn*916*.

### Host-encoded *yqaR* is necessary for *orf17-16*-caused growth arrest

The above results indicate that Orf16, likely in association with Orf17, is interacting with some host cell component to cause cell death. We set out to identify host genes that are required for the cell death caused by Tn*916 orf16* and *orf17* expression. Because expression of *orf17*-*16* causes cell death, we simply isolated suppressor mutations that enable cell survival. We expected to get suppressor mutations that prevent expression of functional *orf17*-*16* from P*xis*. These could include mutations in *orf17*-*16* themselves, or in the regulatory genes (*rapI* and *immA*) needed for derepression of P*xis* through inactivation of the repressor ImmR (Figure 1B). To reduce the frequency of mutations in these genes, we performed this screen with a strain that contained two copies each of *orf17*-*16*, *rapI*, and *immA* (ELC1760). In addition, we included a P*xis*-*lacZ* fusion that would be derepressed similarly to P*xis*-*orf17*-*16*. In this way, we could monitor production of ß-galactosidase to eliminate mutants that prevented P*xis* derepression.

We grew eighteen independent cultures of ELC1760 in defined minimal medium (with 1% arabinose). In early exponential phase, expression of P*xis*-*orf17*-*16* was induced with 1% xylose and cultures were grown overnight (approximately 18 hours). Cultures were diluted and this process was repeated 1-2 times to enrich for suppressor mutants. Cells were plated onto LB agar plates under non-inducing conditions, and candidate mutants were colony-purified and checked for presence of all antibiotic resistance markers. Additionally, we confirmed these isolates properly activated P*xis*-*lacZ* when streaked on LB plates containing X-gal (5-bromo-4-chloro-3- indolyl-β-D-galactopyranoside) and 1% xylose, indicating that the RapI-driven induction of P*xis* was functional (and likely *orf17* and *orf16* were still being expressed). We isolated 18 independent suppressor mutants, one from each of the independent cultures. Genomic DNA from each of these 18 mutants was used for genome resequencing to locate chromosomal mutations.

We found that a single host-encoded gene, *yqaR*, is necessary for the growth arrest caused by Orf17 and Orf16. Fifteen of the 18 mutants were cured of *skin*, a genetic element that interrupts *sigK*, which encodes the mother-cell specific σ^K^ to direct gene expression during sporulation (Kunkel et al., 1990; Stragier et al., 1989). The remaining three mutants each contained a frameshift mutation (either (A)_8→7_ (50/465 nt) or (T)_7→6_ (450/465 nt)) in *yqaR*, a gene in *skin*. The *skin* element is a remnant of a prophage (Takemaru et al., 1995) and contains several homologs of genes in PBSX, a co-resident defective prophage in *B. subtilis* (Krogh et al., 1996). Although *yqaR* is encoded between homologs of a PBSX transcription factor (*yqaQ*) and phage terminase proteins (*yqaST*), there are no homologs of *yqaR* in PBSX (Krogh et al., 1996). Little has been reported about YqaR, although it was identified as a membrane protein found in *B. subtilis* spores (Chen et al., 2019).

We found that deleting either the entire *skin* element or *yqaR* alone eliminated the growth defects and cell killing associated with *orf17*-*16* expression. We generated strains containing a xylose-inducible *orf17*-*16* allele and deleted either the *skin* element (ELC1891) or *yqaR* by introduced a Δ*yqaR*::*cat* allele (ELC1892). These strains were grown in a minimal medium, and growth and viability were monitored by OD and the ability to form CFUs following induction of *orf17*-*16*, as described above. The growth and viability of these strains under inducing conditions was indistinguishable from non-inducing conditions (Figure 5A, 5B), indicating that deletion of *yqaR* or the *skin* element rescues *orf17*-*16* induced killing.

**Figure 5.**
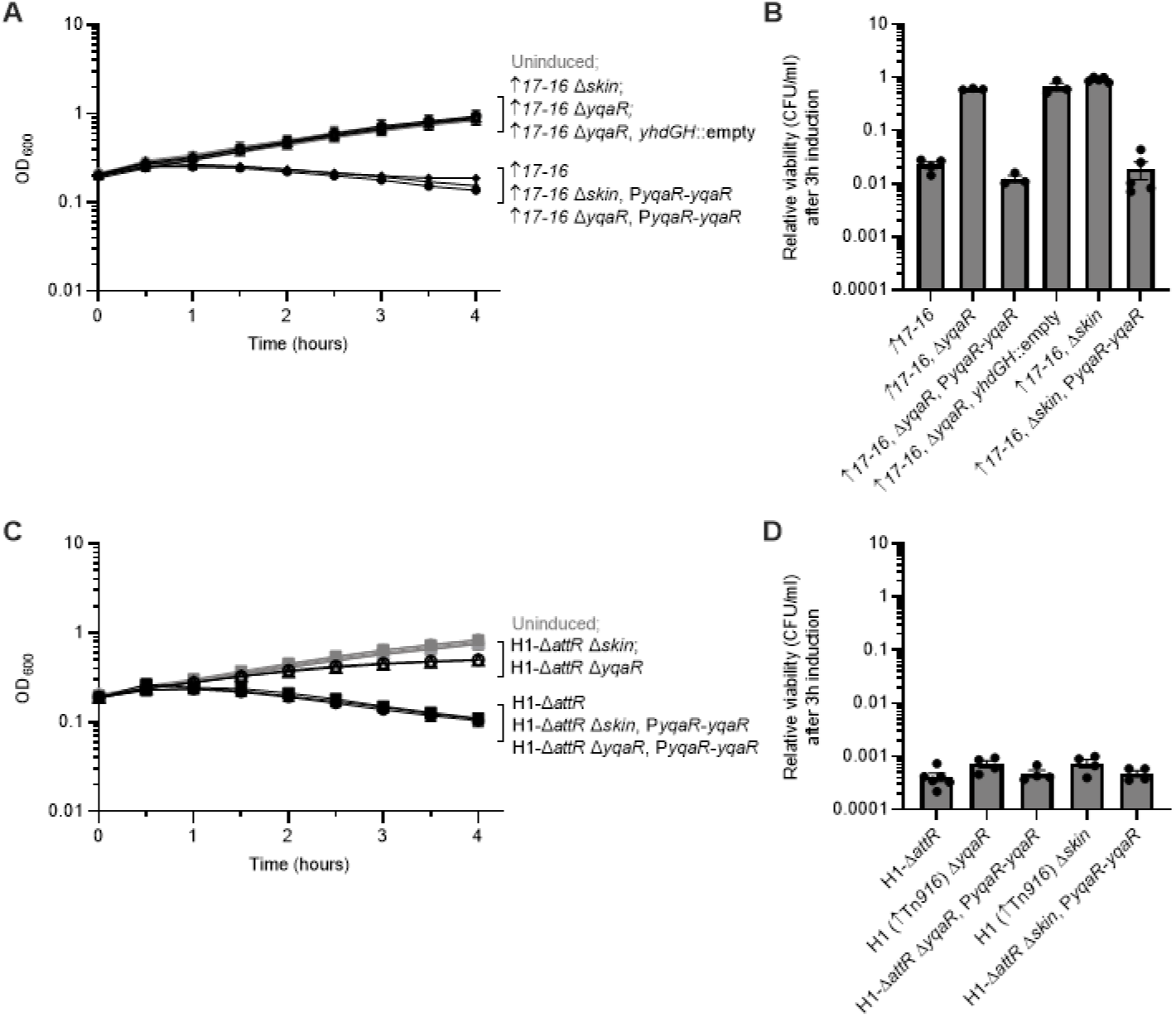
*skin*-encoded *yqaR* is necessary for some Tn*916*-caused growth defects. **A,B)** Strains contained *orf17*-*16* overexpression alleles with the following additional alleles: WT (diamonds, ELC1496), Δ*skin* (open circles, ELC1891), Δ*yqaR* (open upward triangles, ELC1892), Δ*yqaR yhdGH*::empty (open downward triangles, ELC1918), Δ*skin* P*yqaR*-*yqaR* (closed circles ELC1903), and Δ*yqaR* P*yqaR*-*yqaR* (closed upward triangles, ELC1904). **C,D)** Strains contained H1-Δ*attR* with the following additional alleles: WT (squares, ELC1076), Δ*skin* (open circles, ELC1908), Δ*yqaR* (open upward triangles, ELC1856), Δ*skin* P*yqaR*-*yqaR* (closed circles, ELC1909), and Δ*yqaR* P*yqaR*-*yqaR* (closed upward triangles, ELC1911). These strains were grown in minimal arabinose medium to early exponential phase. At time = 0 hours, when cultures were at an OD_600_ ∼0.2, cultures were split into inducing (+1% xylose to stimulate *rapI* expression) and non-inducing conditions. **A,C)** Growth was monitored by OD_600_ for three hours. Black lines indicate growth of the indicated induced cultures; gray lines indicate growth of uninduced cultures. **B,D)** The relative viability of cultures after three hours of element induction was calculated as the number of colony forming units (CFUs) formed by the induced culture, divided by that from the uninduced culture (a value of “1” indicates there is no change in CFUs with induction). Data presented are averages from three or more independent experiments (individual data points shown in (B)), with error bars depicting standard error of the mean. Error bars could not always be depicted in (A) due to the size of each data point.

*yqaR* was the only gene in *skin* needed for the killing caused by expression of *orf17*-*16*. Introduction of a copy of *yqaR*, expressed from its predicted promoter, P*yqaR*, at an ectopic site (*yhdGH*) in the absence of *skin* completely restored the growth defect and cell death caused by expression of *orf17*-*16*. This construct also complemented the *ΔyqaR* mutation. We confirmed that an empty construct inserted at *yhdGH* did not restore the growth arrest and cell killing, indicating that this phenotype was specific to the addition of P*yqaR*-*yqaR*. Altogether, these results indicate that *yqaR* is necessary for effects of *orf17*-*16* on cell growth and viability, and that it is also the only *skin*-encoded gene required for these effects.

### Deleting *yqaR* partially relieves growth defects caused by H1 (Tn*916*) activation

We found that deleting *yqaR* or *skin* in strains containing H1-Δ*attR* partially, but not fully, relieved the growth defects caused by activation of the element. We monitored the growth and viability of strains containing H1-Δ*attR* with the addition of either Δ*skin* or Δ*yqaR*::*cat* (ELC1908 and ELC1856, respectively) following element induction in minimal medium. Deleting *yqaR* or the *skin* element largely improved the OD of the induced cultures until about two hours post-induction, at which point the ODs of these cultures were consistently lower than those of the uninduced cultures (Figure 5C). The introduction of P*yqaR*-*yqaR* into both Δ*skin* or Δ*yqaR*::*cat* strains (ELC1911 and ELC1909, respectively) was sufficient to restore the growth defects caused by H1-Δ*attR* activation. The growth of these strains was indistinguishable from the growth of wild type strains containing H1-Δ*attR*, indicating that *yqaR* is the only *skin*- encoded gene necessary for these detectable growth defects. Consequently, we do not predict that other *skin* gene products are interacting with Tn*916*-encoded proteins.

Deleting *yqaR* or the *skin* element only had a minor effect on CFU recovery following induction of H1-Δ*attR* (Figure 5D). The *ΔyqaR* or Δ*skin* host strains consistently produced slightly more colonies (2-fold greater or less) post-induction than the wild type host strain. The strains in which P*yqaR*-*yqaR* was introduced to complement the deletions behaved indistinguishably from the wild type host strain. These results indicate that *yqaR* does not greatly contribute to the killing effects caused by other Tn*916* genes in the context of H1-Δ*attR*, despite its involvement in the growth defects observed by monitoring OD post-induction.

### Construction of a fluorescent reporter system to visualize Tn*916*-activated cells

These heterologous expression systems revealed that activation of Tn*916* genes negatively impacts population growth and cell viability. Based on these findings, we decided to evaluate the effects of Tn*916* activation on its host cells under its endogenous regulatory system. As Tn*916* is only expressed in ∼1-3% of cells, we sought to do this on a single-cell level.

To monitor potential effects of Tn*916* in its native context on cell growth and viability, we generated a fluorescent reporter to track element activation in single cells. We took advantage of the fact that the DNA processing and conjugation genes in Tn*916* cannot be expressed until the element excises from the chromosome and forms a circular intermediate (Celli and Trieu-Cuot, 1998). Circularization allows promoters on the “right” side of the element to be joined with those genes encoded on the “left” side of the element (Figure 1A). By inserting *gfpmut2* upstream of *orf24* in Tn*916* (Tn*916*-*gfp*), we were able to generate a reporter system in which cells only fluoresce green when the element has been activated and excised (Figure 6A).

**Figure 6.**
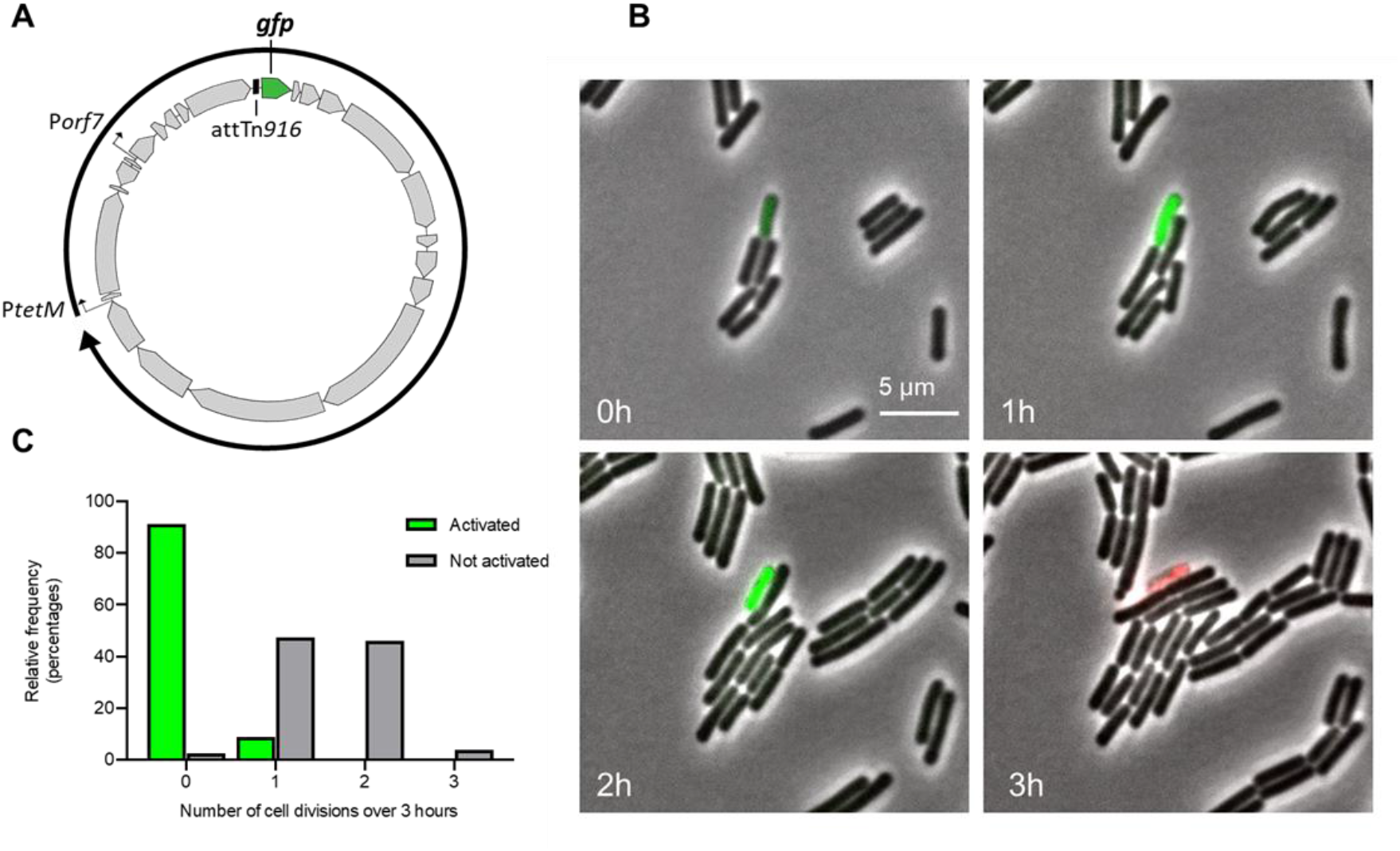
Tn*916*-activated *B. subtilis* cells exhibit growth defects. **A)** The fluorescent reporter system for monitoring Tn*916* excision (Tn*916*-*gfp*). A circular genetic map of Tn*916* is shown, with gray arrows indicating Tn*916* open reading frames, bent arrows representing promoters, and a black box representing the circular attachment site, *att*Tn*916*. P*tetM* and P*orf7* are predicted to drive expression of *orf24*-*orf13* following excision and circularization of the element (reviewed in (Roberts and Mullany, 2009)). *gfpmut2* (shown in green) was inserted upstream of *orf24* such that it will only be expressed following element excision. **B,C)** Cells containing Tn*916*-*gfp* integrated between *yufK*-*yufL* (ELC1458) were grown in minimal glucose medium to late exponential phase with 2.5µg/ml tetracycline to stimulate Tn*916* excision. At time = 0 h, cells were spotted on minimal glucose agarose pads containing 2.5µg/ml tetracycline, 0.1µg/ml propidium iodide, and 0.035µg/ml DAPI. Cells were monitored by phase contrast and fluorescence microscopy for three hours. **B)** A representative set of images from these experiments. GFP (green) was produced in cells in which Tn*916* was activated and excised from the chromosome. Propidium iodide (red) indicates cell death. **C)** Histogram displaying the relative frequency (percentage) of Tn*916*-*gfp* activated cells that underwent the indicated number of cell divisions, compared to non-activated (GFP-negative) cells. DAPI (not shown) was used for monitoring cell division events.

Insertion of *gfp* near the left end of Tn*916*-*gfp* did not abolish excision. We found that after 3 hours of growth in the presence of tetracycline (to stimulate excision), Tn*916-gfp* had excised in ∼0.03% of cells, as measured by qPCR and by counting cells that produced GFP. These results indicated that this reporter is a reliable method to monitor Tn*916* activation in single cells.

### Growth defects caused by Tn*916* activation in *B. subtilis* are observable on the single-cell level

By using this fluorescent reporter system, we found that *B. subtilis* cells in which Tn*916* was activated under its endogenous regulatory system underwent limited to no cell divisions and frequently lysed. Cells containing Tn*916*-*gfp* (ELC1458) were grown in a minimal medium to early exponential phase, then treated with tetracycline to stimulate activation. Three hours later, we visualized cells microscopically, tracking cells that had activated Tn*916*-*gfp* (green, Tn*916*- *gfp* on) and comparing them to cells that had not activated Tn*916*-*gfp* using time-lapse microscopy over the course of three hours.

We tracked 34 cells in which Tn*916*-*gfp* had activated (GFP on) (Figure 6B). Of these 34 cells, 31 (91%) did not undergo any further cell divisions and 3 (9%) divided once (Figure 6C). For comparison, we tracked 76 cells in which Tn*916*-*gfp* had not been activated (GFP negative cells). Of these 76 cells, only 2 (3%) did not divide and 74 (97%) underwent one or more cell divisions.

We also found the activated cells often lysed and died. We used propidium iodide (PI) to monitor the viability of these cells (Boulos et al., 1999). Of the 34 cells that had activated Tn*916*- *gfp*, 91% stained with PI during the course of the experiment (Table 1). In contrast, only 3% of the 76 cells that had not activated Tn*916*-*gfp* were PI-positive or had a PI-positive daughter cell by the end of the time lapse. We did not observe differences in division frequencies or lysis of cells in cases where Tn*916* became activated while on the agarose pad (compared to cells which were activated prior to transferring to the microscopy slide). Results from either case are combined in Figure 6C and Table 1.

**Table 1.** Growth defects caused by Tn916 activation in various host strains^a^

| *B. subtilis* Tn916-*gfp* integration site<sup>b</sup> | Activation state; Mutations | Number cells examined | Cells that divided $\geq 1$ time | PI-stained cells |
| --- | --- | --- | --- | --- |
| *yufK-yufL* | Activated | 34 | 9% | 91% |
|  | Non-activated | 76 | 97% | 3% |
| | Activated; $\Delta orf17-16$ | 36 | 3% | 83% |
| | Activated; $\Delta yqaR$ | 23 | 13% | 65% |
| *ykyB-ykuC* | Activated | 20 | < 5% | 95% |
| *nupQ-maeN* | Activated | 28 | 7% | 79% |

<sup>a</sup>Strains were grown in minimal glucose medium to late exponential phase with 2.5 μg/ml tetracycline to stimulate Tn916 excision. At time = 0 h, cells were spotted on minimal glucose agarose pads containing 2.5 μg/ml tetracycline, 0.1 μg/ml propidium iodide, and 0.035 μg/ml DAPI. Cells were monitored by phase contrast and fluorescence microscopy for three hours. Tn916-*gfp* cells were monitored in both activated (GFP-positive) and non-activated (GFP-negative) states and the number of cell divisions was counted. Cells were counted as dead if either they or any daughter cells turned PI-positive during the 3-hour time lapse.
<sup>b</sup>Host strains examined: Tn916-*gfp* integrated in 3 different chromosomal locations: between *yufK-yufL* (ELC1458), *ykuC-ykyB* (LKM20), and *nupQ-maeN* (LKM18), and two mutant strains with Tn916-*gfp* integrated between *yufK-yufL* with the indicated deletions: $\Delta orf17-16$ (ELC1512) and $\Delta yqaR$ (ELC1857).

We were concerned that the growth defect and loss of viability that we observed might be due to treatment with tetracycline. This was not the case. Neither growth arrest nor cell death were dependent on the presence of tetracycline. Tetracycline modestly enhances, but is not necessary for Tn*916* activation (Celli and Trieu-Cuot, 1998; Manganelli et al., 1995; Showsh and Andrews, 1992; Su et al., 1992; Wright and Grossman, 2016). To confirm that the presence of the antibiotic was not impacting the growth of activated cells, we monitored the growth and viability of Tn*916* activated cells without the addition of tetracycline in conditions otherwise identical to those described above. We identified 20 cells that had activated Tn*916*-*gfp*. Of these 20 cells, 19 (95%) did not divide, and all 20 (100%) lysed during the course of the experiment (Table 1). These results are indistinguishable from those observed with tetracycline added. Because tetracycline modestly enhances Tn*916* activation, consequently increasing the number of activated cells to track, it was included for the rest of the experiments.

We found that the growth and viability defects caused by excision of Tn*916*-*gfp* from the insertion site between *yufK* and *yufL* were not unique to the particular Tn*916* genomic insertion site. Tn*916* can insert into many sites in AT-rich regions in the bacterial chromosome (Clewell et al., 1988; Cookson et al., 2011; Mullany et al., 2012; Scott et al., 1994)). Therefore, we monitored the growth of cells in which Tn*916*-*gfp* was located at two different locations on the chromosome, one inserted between *ykuC* and *ykyB* (LKM20) and one between *nupQ* and *maeN* (LKM18). We observed similar results of limited divisions and frequent lysis for these two insertion sites compared to the insertion between *yufK* and *yufL* (Table 1), indicating these results were not dependent on the integration site of Tn*916*. Interestingly, the site between *ykuC* and *ykyB* is as a “hot spot” for Tn*916* integration in the *B. subtilis* chromosome indicating that these growth defects occur upon excision from a preferred integration site in the chromosome.

Together, these results indicate that cells in which Tn*916* became activated were unable to divide and lost viability. These results are consistent with those we observed on the population-level using the ICE*Bs1*-Tn*916* hybrid (H1), that activates Tn*916* genes in the majority of cells in a population.

## Host cells lacking *orf17***-***16* or *yqaR* exhibit growth defects upon element activation

Because deleting *orf17*-*16* or *yqaR* improved the growth and delayed the decrease in viability in the population-based assays expressing Tn*916* genes in the majority of cells, we reasoned we might see similar improvements on the single-cell level while monitoring the growth and viability of Tn*916* host cells with deletions of *orf17*-*16* or *yqaR*. However, deletions of these genes did not rescue growth and viability of Tn*916* activated cells. Under conditions identical to those described above, we monitored the number of divisions and viability of Δ*orf17*-*16* and Δ*yqaR* Tn*916* host cells (ELC1512, ELC1857, respectively). These deletions did not drastically improve the number of cell divisions of the activated host cells (Table 1). However, the Δ*orf17*- *16* and Δ*yqaR* host cells did have lower percentages of cells become PI-positive compared to WT hosts with Tn*916* at the same chromosomal locus during the 3-hour time lapse (83% and 65%, respectively, compared to 91% for WT). These microscopy assays may lack the resolution necessary to detect significant differences in growth of mutant backgrounds and are limited by the necessarily small sample sizes. Overall, these results indicated that these mutations did not enable restored growth or viability of activated cells. These results further support the notion that Tn*916* possesses multiple mechanisms to manipulate host cell growth and viability beyond the relationship between Orf17-16 and YqaR.

### Deleting *yqaR* in Tn*916* host cells increases conjugation efficiency

We wondered if these unexpected interactions between Tn*916* and *B. subtilis* host cells would have any effect on how well the element could move through a population. Perhaps these deleterious phenotypes would limit the ability of Tn*916* to efficiently move from cell-to-cell. Alternatively, these phenotypes could indicate that Tn*916* is taking advantage of its host to facilitate its likely energetically-costly transfer events (Reinhard et al., 2013; Ryan et al., 2016).

We found that *yqaR* null donors mated more efficiently than their wild-type counterparts. We evaluated the impact of *yqaR* presence in donor and recipient cells on transfer efficiencies; we could not test the effects *orf17* or *orf16* deletions because the proteins encoded are required for Tn*916* conjugation. Donor strains (± *yqaR*) containing a copy of Tn*916* between *yufK* and *yufL* and recipient strains (± *yqaR*) were grown in LB medium to early exponential phase. At this point, Tn*916* activation was stimulated in donor cultures through the addition of 2.5 µg/ml tetracycline. After a one-hour induction period, donor and recipient cultures were mixed, filtered, and applied to a solid surface for mating. After one hour, cells were resuspended and the number of transconjugant CFUs (recipients that acquired Tn*916*) was determined. Conjugation efficiencies were calculated by normalizing the number of transconjugant CFUs to the number of donor CFUs; conjugation efficiencies of matings using Δ*yqaR* strains were normalized to those of matings using WT strains that were conducted in parallel for each experimental replicate.

Δ*yqaR* donors (ELC1851) mated approximately 10-fold more efficiently than their *yqaR*+ (CMJ253) counterparts (Figure 7A). However, deleting *yqaR* in recipient cells (ELC1854) did not lead to detectable differences in mating efficiencies compared to *yqaR*+ recipients (ELC301), indicating that the *yqaR* effects are specific to donor cells. In parallel experiments, we confirmed that no other *skin*-encoded genes impact Tn*916* transfer efficiency. Δ*skin* donor cells (ELC1846) exhibited a similar improvement in mating efficiency to Δ*yqaR* donor cells. Additionally, complementing *yqaR* under its native promoter at an ectopic locus (*yhdGH*) restored Tn*916* mating efficiencies to WT levels. These results indicate that the presence of *yqaR* negatively affects the transfer efficiency of Tn*916*.

**Figure 7.**
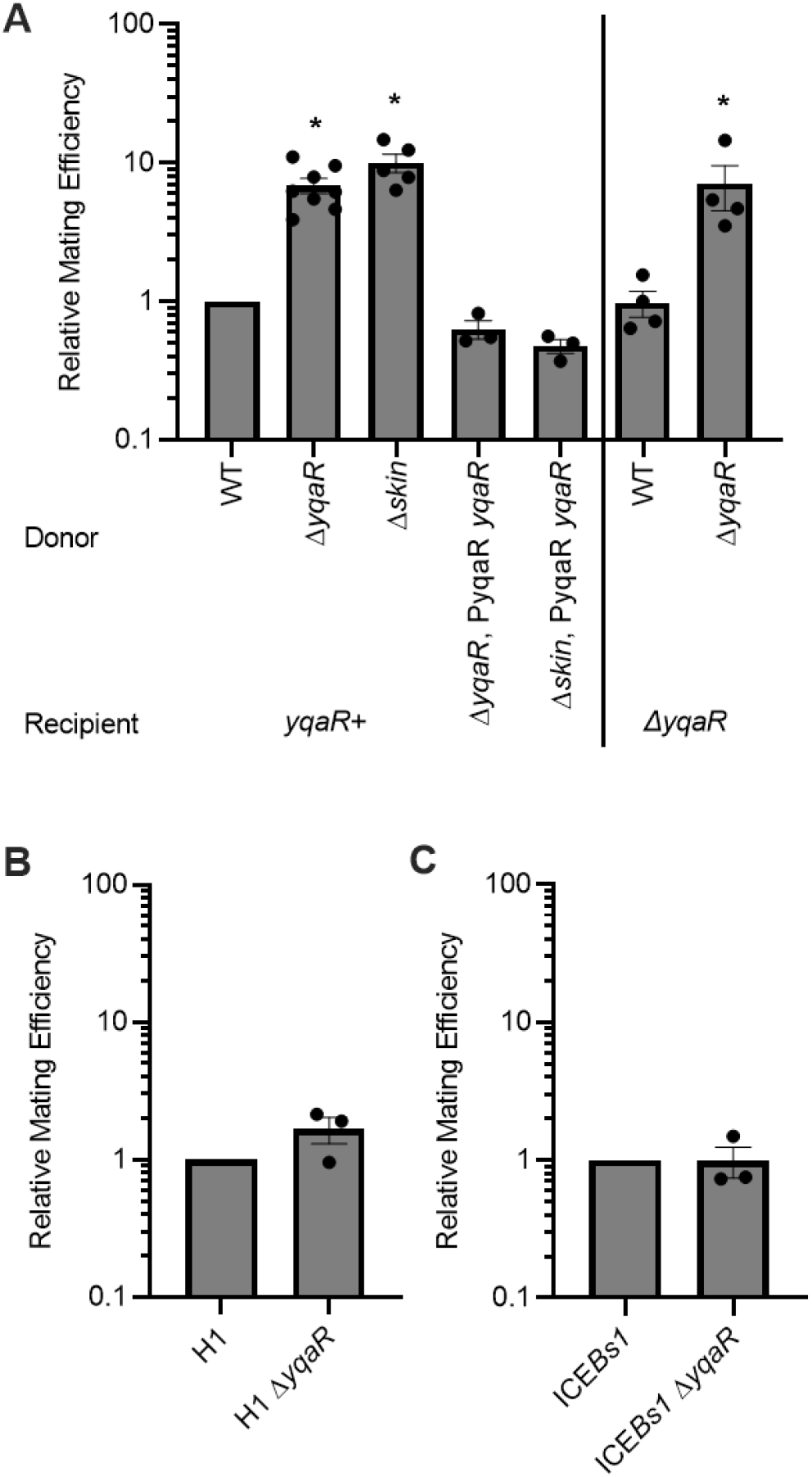
Deleting *yqaR* in Tn*916* donors increases mating efficiency. **A)** Tn*916* donors: WT (CMJ253), Δ*yqaR* (1851), Δ*skin* (ELC1846), Δ*yqaR* P*yqaR*-*yqaR* (ELC1922), Δ*skin* P*yqaR*-*yqaR* (ELC1923) were grown to early-exponential phase in LB medium, Tn*916* activation was stimulated with 2.5 µg/ml tetracycline for one hour, and then donors were mated with *yqaR*+ (ELC301) or Δ*yqaR* (ELC1854) recipients. Conjugation efficiencies (the number of transconjugants produced divided by the number of donors applied to mating) were normalized to those calculated for WT Tn*916* donors mated into *yqaR+* recipients, which were completed in parallel for each experimental replicate. **B,C)** Donor strains containing H1 (ELC1213), H1 Δ*yqaR* (ELC1843), ICE*Bs1* (JMA168), or ICE*Bs1* Δ*yqaR* (ELC1844) were grown to early-exponential phase in LB, ICE activation was stimulated with 1 mM IPTG for one hour, and then donors were mated with *yqaR*+ (ELC301) recipients. Conjugation efficiencies of Δ*yqaR* matings were normalized to *yqaR*+ donor matings conducted in parallel. Typical conjugation efficiencies in these experiments were as follows: Tn*916* ∼0.0005%, H1 ∼0.5%, ICE*Bs1* ∼1%. Data shown are an average of three or more independent experiments. Error bars represent standard error of the mean.

We found that the effects of *yqaR* on Tn*916* conjugation are downstream of the activation step of the element’s lifecycle. Deleting *yqaR* did not alter the activation frequency of Tn*916*. Tn*916* had excised (and was therefore activated) in 0.79 ± 0.13% of Δ*yqaR* donors compared to 0.83 ± 0.13% of WT Tn*916* donors immediately prior to the start of the mating.

Deleting *yqaR* had less impact on the transfer efficiency of the hybrid conjugative element (ICE*Bs1*-Tn*916*)-H1. Donor strains (± *yqaR*) containing a copy of H1 and an IPTG-inducible copy of the activator *rapI* were grown in LB medium to early exponential phase, induced with 1mM IPTG for one hour, and mated with a recipient culture in identical conditions to those described above. H1 Δ*yqaR* donors (ELC1843) mated only ∼2-fold more efficiently than *yqaR*+ donors (ELC1213) (Figure 7B). This result indicates that *yqaR* has less of an impact on conjugative transfer of H1 than Tn*916*. It is possible that the increased activation frequency of H1 and consequently increased transfer efficiencies is masking the effects of *yqaR* on mating efficiencies.

Because ICE*Bs1* activation does not cause growth defects like those elicited by Tn*916*, we did not expect *yqaR* to influence ICE*Bs1* mating efficiencies. Indeed, under identical mating conditions to those described above, Δ*yqaR* ICE*Bs1* donors (ELC1844) did not exhibit altered mating efficiencies compared to *yqaR*+ ICE*Bs1* donors (MMB766) (Figure 7C). This result confirms that *yqaR* is specifically interacting with Tn*916*-encoded genes, and not broadly impacting the transfer of conjugative elements.

Together, these results indicate that the YqaR-dependent growth defects caused by Tn*916* genes are not beneficial for efficient element transfer. Instead, it appears that *B. subtilis* encodes a mechanism to limit the spread of Tn*916* through a population of cells. However, these results do not elucidate any effects that other Tn*916*-mediated growth defects may have on transfer efficiency.

### Tn*916* activation causes growth defects in an *Enterococcus faecalis* strain

Whereas *B. subtilis* is a convenient host for analyzing Tn*916* (e.g., (Christie et al., 1987; Ivins et al., 1988; Mullany et al., 1990; Roberts et al., 2003; Scott et al., 1988; Wright and Grossman, 2016)), it is not a natural host. Furthermore, homologs of YqaR are not found in any of its natural hosts (*Enterococcus*, *Streptococcus*, *Staphylococcus*, and *Clostridium* species). Therefore, we wondered if this killing elicited by activated Tn*916* is specific to *B. subtilis* or might also occur in a natural host.

We found that Tn*916* activation-related growth defects and death also occur in *Enterococcus faecalis*, a natural Tn*916* host. We monitored the effects of Tn*916* activation in *Enterococcus faecalis*, the first-discovered host species of Tn*916* (Franke and Clewell, 1981a, 1981b) using the same fluorescent reporter system described above. Tn*916*-*gfp* was mated into *E. faecalis* (ATCC 19433) and two independent transconjugants were isolated to evaluate the effects of activation (excision) of Tn*916*-*gfp* from different chromosomal integration sites. The integration sites for two isolates (strains ELC1531 and ELC1529) were each in a different chromosomal location, and strain ELC1529 had two distinct chromosomal copies of Tn*916*-*gfp* (Materials and Methods). Each strain was grown to early exponential phase in a supplemented M9 medium (Materials and Methods). Tetracycline (2.5 µg/ml) was then added to stimulate excision of the element. After an hour of growth with tetracycline, cells were visualized for two hours using time-lapse microscopy (Figure 8A).

**Figure 8.**
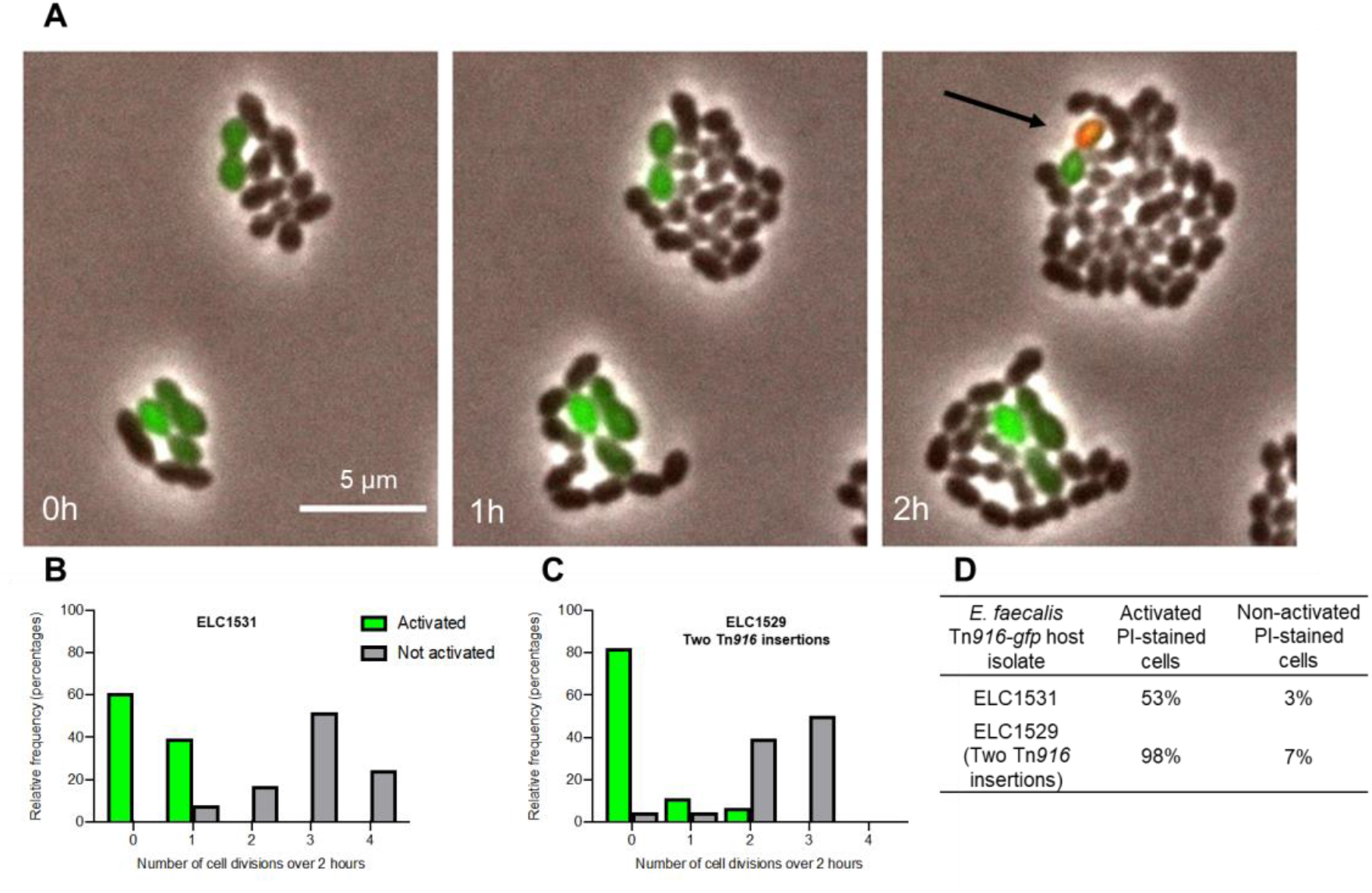
Tn*916*-activated *E. faecalis* cells exhibit growth defects. **A-D)** Two separate isolates of *E. faecalis* (ATCC 19433) containing Tn*916*-*gfp* were used to monitor effects of Tn*916* activation. Tn*916*-*gfp* was inserted in the following sites (integration site sequences in Materials and Methods): ELC1531 has one copy between ABC_ACP; ELC1529 has two copies: between *citG_*SP and HP_CSP. Cells were grown in M9 medium to late exponential phase with 2.5µg/ml tetracycline to stimulate Tn*916* excision. At time = 0 h, cells were spotted on M9 medium agarose pads containing 2.5µg/ml tetracycline, 0.1µg/ml propidium iodide, and 0.5µg/ml DAPI. Cells were monitored by phase contrast and fluorescence microscopy for two hours. **A)** A representative set of images monitoring ELC1531 cells with an activated copy of Tn*916*-*gfp* (GFP-positive). Similar results were observed with ELC1529. The black arrow in the final frame indicates a PI-stained, GFP-positive cell (appears reddish-yellow). **B,C)** Histograms displaying the relative frequency (percentage) of Tn*916*-*gfp* activated cells that underwent the indicated number of cell divisions, compared to non-activated (GFP-negative) cells for each isolate. For ELC1531 and ELC1529: 66 and 45 activated and non-activated cells were monitored for each, respectively. **D)** The frequency of cells that became PI-stained (indicating cell death) or had a daughter cell become PI-stained was determined for both activated and non-activated cells during the 2- hour time lapse. ABC = ABC transporter; ACP = acyl carrier protein; SP = surface protein; CSP = cold shock protein (Annotations based on *E. faecalis* KB1 annotations, Accession number: CP015410.1).

Activated cells from both isolates exhibited a decrease in cell divisions and were stained with PI more frequently than neighboring non-activated cells (Figure 8B-D). From ELC1531, 66 cells that were producing GFP (had an excised Tn*916-gfp*) were tracked: 26 cells (39%) underwent one division and 35 (53%) appeared to lose viability based on staining with PI during the time lapse. In contrast, of the 66 non-activated cells that were tracked, 100% underwent one or more divisions and only 2 cells (3%) became PI-positive or had progeny that became PI-positive. From strain ELC1529, 45 cells were tracked: 8 cells (18%) underwent one or more divisions and 98% appeared to lose viability based on staining with PI during the time lapse; of the 45 non-activated cells tracked, 43 cells (96%) underwent one or more divisions and only 3 (7%) became PI-positive. It is noteworthy to point out that *E. faecalis* does not grow on the minimal medium used for *B. subtilis* microscopy; instead, we used a rich medium to grow and visualize *E. faecalis* during live cell imaging, contributing to a faster growth rate than that for *B. subtilis*. Thus, we cannot directly compare the number of divisions of activated Tn*916* between *B. subtilis* and *E. faecalis* host cells.

The growth defects were most pronounced in ELC1529, which has two copies of Tn*916* in the chromosome. We predict that having two copies of Tn*916* exacerbates the growth phenotypes following Tn*916* activation. It is likely that the activation and excision of one copy of Tn*916* leads to activation and excision of the second copy (Flannagan and Clewell, 1991; Lunde et al., 2019; Manganelli et al., 1995). Thus, having two copies of Tn*916* in a host cell likely leads to higher expression levels of detrimental gene(s), which could be deleterious for the host. This result is interesting given that Tn*916* does not possess any known mechanisms to prevent acquisition of multiple copies in a single host, such as exclusion systems that are used by conjugative plasmids and some ICEs (Avello et al., 2019; Garcillán-Barcia and de la Cruz, 2008; Marrero and Waldor, 2005).

Although we did not identify a homolog of YqaR in *E. faecalis*, we confirmed that the mechanism of Tn*916* deleterious interactions with *E. faecalis* host cells was independent of *orf17*-*orf16*. We mated a Tn*916* (*orf17*-*orf16*) mutant from a *B. subtilis* donor into *E. faecalis* (ELC1696; Materials and Methods). We monitored the growth of 23 activated cells in conditions identical to those described above. Of these, 12 cells (52%) underwent one or more division and 11 (23%) appeared to lose viability based on PI staining. These results are similar to those observed for ELC1531, indicating that deleting *orf17*-*orf16* from Tn*916* did not detectably improve growth of host cells following activation. It is possible that these experiments do not possess the resolution necessary to detect growth improvements, as noted for *B. subtilis* microscopy experiments described above. However, these results highlight that Tn*916* possesses a mechanism independent of *orf17*-*orf16* to modulate host growth in viability in multiple host species, not just *B. subtilis*. In conjunction with results reported above, Tn*916* appears to encode multiple mechanisms to influence host growth, and at least some of these mechanisms may function similarly in various hosts.

## DISCUSSION

The experiments described here demonstrate that activation of Tn*916* causes a growth arrest and cell death. We suspect that these effects were previously undetected due to the low activation frequency of the element. We were able to detect the growth arrest and cell death caused by Tn*916* by studying a hybrid ICE that can be activated in a large fraction of cells in a population, and by analyzing the low activation frequency Tn*916* in single cells using a fluorescent reporter.

Two previous reports contained results indicating that when activated, Tn*916* was deleterious to the host cell. We previously noted a decline in the percentage of cells in which Tn*916* had excised, leading us to speculate that Tn*916* could have some deleterious effect on cell growth (Wright and Grossman, 2016). Additionally, a previous report found that *E. faecalis* host cells containing a Tn*916* mutant with increased excision frequencies (due to mutations in the regulatory region upstream of *tetM*) had decreased fitness relative to cells containing wild type Tn*916* (Beabout et al., 2015). The authors hypothesized that the decreased fitness was due to increased production of TetM, possibly slowing protein production, and an additional cost of the hyper-conjugative phenotype of the mutant (Beabout et al., 2015). We suspect that in both these studies (Beabout et al., 2015; Wright and Grossman, 2016), the decreased fitness was due to the growth arrest and cell killing described here.

### Interactions between a conjugative element (Tn*916*) and a defective prophage-like element (*skin*)

We found that multiple Tn*916*-encoded genes contribute to the growth arrest and cell death phenotype upon element activation. Growth arrest and cell death caused by Orf17-16 is dependent on *yqaR*, a gene found within the defective phage-like element *skin*. Loss of *yqaR* (or *skin*) leads to an increase in Tn*916* conjugation, indicating that one function of *yqaR* might be to limit activity and spread of this conjugative element. These interactions are reminiscent of an abortive infection mechanism (Labrie et al., 2010). An intriguing comparison is the abortive infection mechanism encoded by ICE*Bs1*, in which *spbK* (from ICE*Bs1*) protects host cells from predation by the co-resident prophage SPß (Johnson et al., 2022). However, in the case of Tn*916* and the defective prophage-like element *skin*, it is the prophage that appears to be limiting spread of the conjugative element. Following the abortive infection analogy, the ability of a host cell to limit spread of Tn*916* might protect neighboring cells from killing by Tn*916*. Alternatively, host cells may limit the spread of Tn*916* to limit the sharing of *tetM* and outcompete neighboring cells in the presence of tetracycline. Because Tn*916* is only activated in a small fraction of cells in a population, this model could be feasible. The interactions (direct or indirect) between the *yqaR* and *orf17*-*16* gene products represents a type of interaction and perhaps competition between mobile genetic elements that share a bacterial host.

### Multiple mechanisms by which Tn*916* causes growth arrest and cell death

Beyond the growth arrest and killing mediated by *yqaR* and *orf17*-*16*, Tn*916* has other genes that cause host cell death. In the absence of *yqaR*, there is growth arrest and cell death when Tn*916* is activated in *B. subtilis*. In addition, growth arrest and cell death also occur in *E. faecalis*, which has no recognizable homologs of *yqaR*. It is possible that there are functional analogs of YqaR, but we favor a model in which Tn*916* influences cell growth and viability through other pathways, both in its natural host *E. faecalis* and in *B. subtilis*. We suspect that similar processes occur in other natural hosts, and that close relatives of Tn*916* are likely to cause similar phenotypes.

### Cell fate and the spread of integrative and conjugative elements

A major question arising from our findings centers around the fate of the transconjugant cells that acquire a copy of Tn*916*. During conjugation, a linear single-stranded copy of Tn*916* is transferred from donor to recipient. Once in the recipient, the DNA recircularizes and is replicated to form a dsDNA circle, which is the substrate for integration (Johnson and Grossman, 2015). The Tn*916* genes are presumably expressed from the dsDNA; in particular, the integrase needs to be made. Based on our results, we expect that expression of Tn*916* genes would be detrimental to the nascent transconjugants. However, it is clear that Tn*916* is able to successfully transfer and produce viable transconjugants, indicating that at least some fraction of nascent transconjugants are able to survive. Perhaps Tn*916* is able to integrate and thus halt expression of its detrimental genes in a short time scale that does not compromise the viability of its host cell. The initial acquisition of a conjugative element can be costly to host cell growth, and such a phenotype would not be unique to Tn*916*; previous reports have demonstrated the costs associated with conjugative element acquisition (e.g. Dahlberg and Chao, 2003; Doyle et al., 2007; Nguyen et al., 2011; Prensky et al., 2021). Future studies may explore the mechanisms of cell survival in transconjugants.

Similar questions apply to other ICEs. For example, ICE*clc* from *Pseudomonas* species is activated stochastically in 3-5% of cells in a population, and these cells differentiate into a “transfer competent” state that is characterized by slow growth, decreased viability, and the ability to transfer the element efficiently (Delavat et al., 2016; Reinhard et al., 2013). The genes required for the decreased cell growth and viability do not encode components of the conjugation machinery, but their loss causes a decrease in conjugation efficiency, indicating that the differentiated state is somehow important for efficient transfer of ICE*clc* (Reinhard et al., 2013). This is in contrast to the situation with Tn*916*: at least some of the Tn*916* genes that contribute to the growth arrest and cell death encode proteins that are part of the conjugation machinery.

More similarly to Tn*916*, an essential component of the conjugation machinery encoded by the ICE R391, originally isolated from *Providencia rettgeri*, permeabilizes the host cell membrane, causing death (Armshaw and Pembroke, 2013a, 2013b). This killing is proposed to function as a back-up mechanism for spread of the element through a population (Armshaw and Pembroke, 2015; Ryan et al., 2016). However, the mechanisms by which Tn*916* causes growth and viability defects appears to be different. First, the protein in R391 responsible for these phenotypes is not related to any of those encoded by Tn*916*. Second, for Tn*916*, partial alleviation of the growth and viability defects leads to an increase in transfer. Thus, for R391 and ICE*clc*, the growth arrest and decreased viability stimulate transfer whereas for Tn*916*, they inhibit transfer. These differences highlight how various ICEs have evolved multiple and contrasting mechanisms to impact host growth and viability.

We suspect that other ICEs have similarly complex impacts on their host cells. However, since most ICEs are activated in only a relatively small fraction of cells in a population, these effects are difficult to observe. The ability to activate an ICE in a large fraction of cells and to visualize and analyze individual cells that contain an active ICE should reveal many of the complex interactions that occur between an ICE, a host cell, and other horizontally acquired elements.

## MATERIALS AND METHODS

### Media and growth conditions

*B. subtilis* cells were grown shaking at 37°C in either LB medium or MOPS (morpholinepropanesulfonic acid)-buffered 1XS7_50_ defined minimal medium (Jaacks et al., 1989) containing 0.1% glutamate, required amino acids (40 μg/ml phenylalanine and 40 μg/ml tryptophan) and either glucose or arabinose (1% (w/v)) as a carbon source or on LB plates containing 1.5% agar.

*E. faecalis* cells were grown shaking at 37°C either in an M9 medium, consisting of 1X M9 salts supplemented with 0.3% yeast extract, 1% casamino acids, 3.6% glucose, 0.012% MgSO_4_, and 0.0011% CaCl_2_ (Breuer et al., 2017; Dunny and Clewell, 1975) or in BHI medium. *Escherichia coli* cells were grown shaking at 37°C in LB medium for routine strain constructions. As appropriate, antibiotics were used in standard concentrations (Harwood and Cutting, 1990): 5 μg/ml kanamycin, 12.5 μg/ml tetracycline, and 100 μg/ml streptomycin for selection on solid media.

### Strains, alleles, and plasmids

*E. coli* strain AG1111 (MC1061 F’ *lacI*^q^ *lacZ*M15 Tn*10*) was used for plasmid construction. *Enterococcus faecalis* (ATCC-19433) was used as a host strain for Tn*916*-*gfp*.

*Bacillus subtilis* strains (Table 2), except BS49, were derived from JH642, contain the *trpC2 pheA1* alleles (Perego et al., 1988; Smith et al., 2014), and were made by natural transformation (Harwood and Cutting, 1990). The construction of newly reported alleles is summarized below.

**Table 2.**
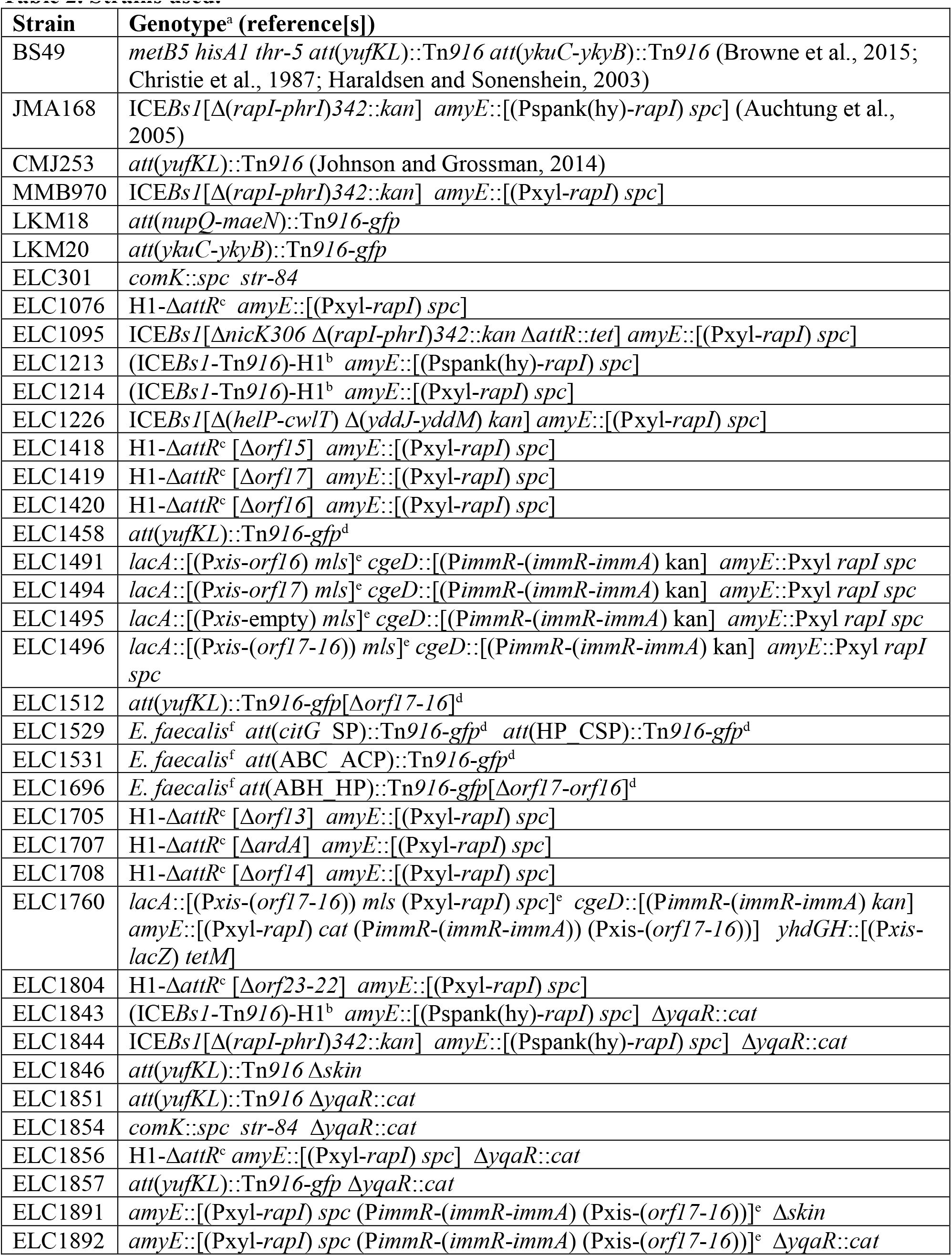
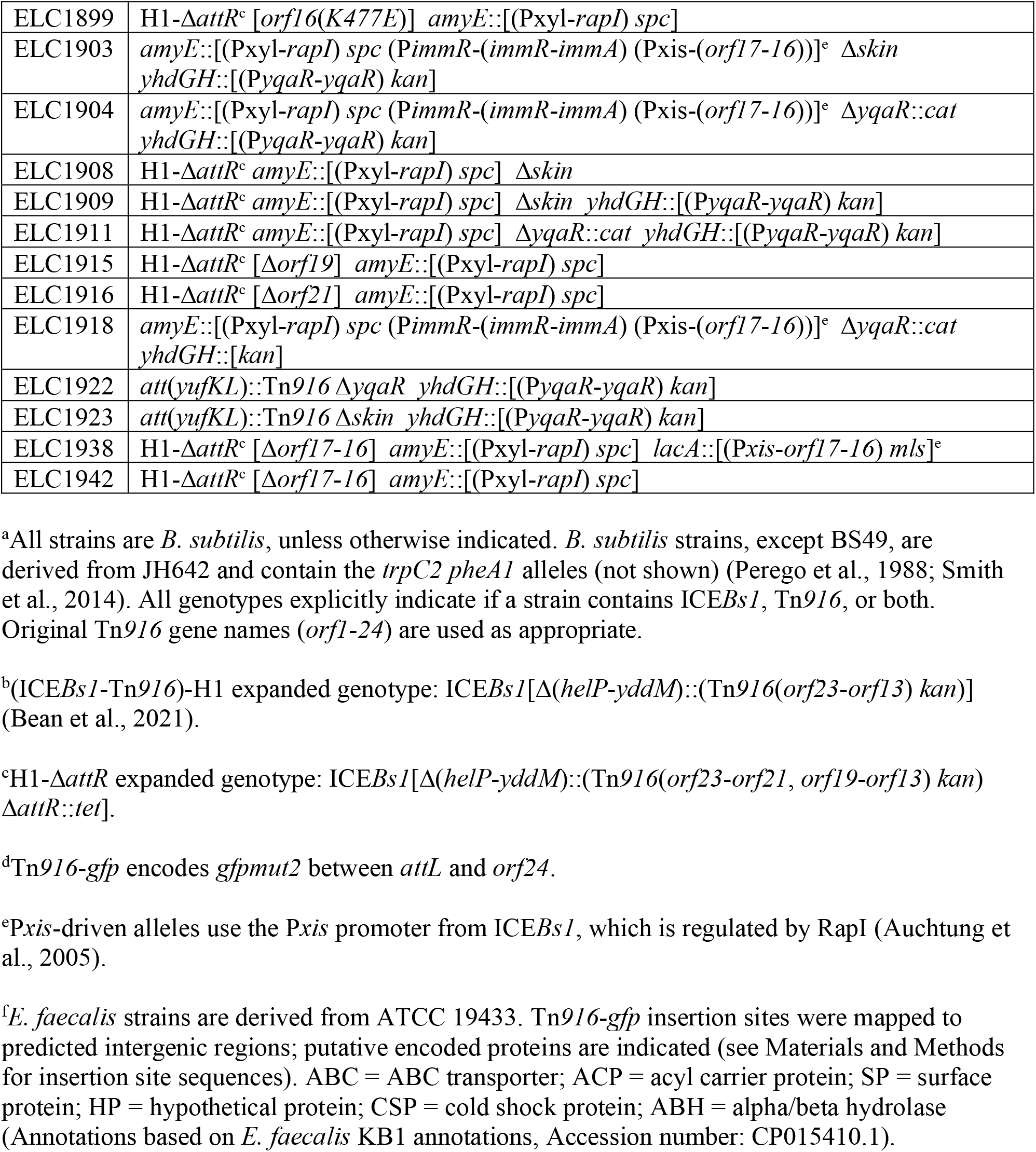
Strains used.

Tn*916* host strain CMJ253 contains a single copy of Tn*916* between *yufK* and *yufL* (Johnson and Grossman, 2014). It was generated by natural transformation of JMA222, a JH642-derived strain that was cured of ICE*Bs1* (Auchtung et al., 2005), with genomic DNA from BS49 (Browne et al., 2015; Christie et al., 1987; Haraldsen and Sonenshein, 2003), selecting for tetracycline resistance from Tn*916* acquisition, as previously described (Johnson and Grossman, 2014; Wright and Grossman, 2016).

The construction of (ICE*Bs1*-Tn*916*)-H1, a hybrid integrative and conjugative element, was previously described (Bean et al., 2021). Essentially, we replaced the DNA processing and conjugation genes of ICE*Bs1* (*helP*-*cwlT*) with those of Tn*916* (*orf23-orf13*) (Figure 1). In doing so, the Tn*916* genes are controlled by the P*xis* promoter from ICE*Bs1* (regulated by ImmR, ImmA, and RapI), and use the integration/excision components of ICE*Bs1* (Int, Xis). Therefore, we can activate Tn*916* gene expression in a greater proportion of cells than in wild type Tn*916* through overproduction of the ICE*Bs1*-encoded activator protein RapI (Bean et al., 2021).

The construction of the Δ(*rapI*-*phrI*)*342*::*kan* allele used to track transfer of ICE*Bs1* during matings was described previously (Auchtung et al., 2005). The Δ*attR*::*tet* allele (Lee et al., 2007) used to prevent element excision from the chromosome, and Δ*nicK* (Lee and Grossman, 2007), Δ*orf20* (Wright and Grossman, 2016) were all previously described.

ICE*Bs1*, (ICE*Bs1*-Tn*916*)-H1, and complementation constructs were under the regulatory control of P*xis* (of ICE*Bs1*) and were activated by overexpression of *rapI* using either a xylose-inducible copy of *rapI* ((*amyE*::[(Pxyl-*rapI*) *spc*]) (Berkmen et al., 2010) or an isopropyl-β-D- thiogalactopyranoside (IPTG)-inducible copy of *rapI* (*amyE*::[(Pspank(hy)-*rapI*) *spc*]) (Auchtung et al., 2005). ICE-cured strains, which did not contain a copy of the P*xis* repressor, *immR*, contained an ectopic allele for normal P*xis* regulation *cgeD*::[(P*immR*-(*immR*-*immA*) kan] (Auchtung et al., 2007).

*B. subtilis* strains cured of ICE*Bs1* with the streptomycin resistance allele (*str*-*84*) were previously described (Auchtung et al., 2005; Luttinger et al., 1996). The Δ*comK*::*spc* allele in ELC301 replaced most of the *comK* open reading frame from 47bp upstream of *comK* to 19bp upstream of its stop codon with the spectinomycin resistance cassette from pUS19. The *spc* marker was fused with up- and downstream homology regions via isothermal assembly (Gibson et al., 2009) and used for transformation.

Unmarked deletions were generated for Tn*916* genes *orf23*-*orf13* in the context of the hybrid element. Briefly, flanking homology regions were amplified for each deletion and inserted by isothermal assembly into pCAL1422, a plasmid containing *E. coli lacZ* and *cat* in the backbone, cut with EcoRI and BamHI (Thomas et al., 2013). The resulting plasmids were used to transform an appropriate *B. subtilis* strain, selecting for single crossover of the plasmid into the chromosome (chloramphenicol resistant). Transformants were screened for loss of *lacZ* and checked by PCR for the desired deletion. The deletion boundaries are described below.

The Δ*orf23*-*22* deletion starts immediately after the *orf24* stop codon through the *orf22* stop codon. The *Δorf21* deletion, removes the first 1272 bp of *orf21* and leaves the remaining 114 bp (and *oriT*) intact. The Δ*orf19* deletion starts 5bp upstream of *orf19* and ends 15bp upstream of *ardA*, likely abolishing *sso916* which is encoded between *orf19* and *ardA* (Wright and Grossman, 2016). The Δ*ardA* deletion removes most of the *ardA* ORF, leaving the final 26bp intact (to leave a previously misannotated *orf17* start codon intact). Of note, we found that *orf17* actually begins 88 bp downstream of *ardA* (it was previously predicted to overlap with the last 26bp of *ardA*; the actual start codon was previously predicted to be Met39). The misannotated start site lacked an obvious ribosome binding site; we found that an ectoptic expression allele using the *orf17* “downstream” start site was able to restore mating for a donor strain containing (ICE*Bs1*-Tn*916*)-H1(Δ*orf17*), which could not detectably mate. The Δ*of17* deletion starts 78bp upstream of the *orf17* start codon and leaves the last 14 codons intact. The Δ*orf16* deletion removes codons 10-804 (of 815 total). The Δ*orf15* deletion was designed to remove codons 6- 716 (of 754 total), based on the sequence for Tn*916* in *Enterococcus faecalis* DS16 (Genbank U09422.1). However, *orf15* in Tn*916* from BS49 contains a cytosine insertion resulting in a 725- amino acid protein (Browne et al., 2015). This frameshift was removed in the deletion, allowing codons 1-5 to be fused with the originally annotated codons 716-754. The Δ*orf14* deletion removes codons 32-325 (of 333 total).

A similar strategy was used to generate an unmarked missense mutation in the predicted Walker A motif of *orf16* to eliminate nucleotide binding. *Orf16*(*K477E*) was generated via isothermal assembly of homology arms into pCAL1422. The resulting plasmid (pELC1295) was used to acquire the desired point mutant, by transforming into *B. subtilis* and subsequently screening for the desired mutant. A similar approach was used to generate *conE* (*K476E*) in ICE*Bs1* (Berkmen et al., 2010). A donor strain containing (ICE*Bs1*-Tn*916*)-H1 (*orf16*(*477E*)) did not detectably transfer in standard mating assays, confirming the indicated missense mutation disrupted typical Orf16 mating functions.

Ectopic expression alleles controlled by P*xis* were designed at *lacA*, as previously described (Jones et al., 2021), to test the sufficiency of Tn*916* genes to cause growth defects. Briefly, *orf16*, *orf17*, or *orf17*-*16* together were fused with P*xis*, an MLS resistance cassette, and up- and downstream homology arms via isothermal assembly and transformed into *B. subtilis*, selecting for acquisition of MLS resistance. A *lacA*::[(P*xis*-empty) *mls*] control allele did not insert an ORF downstream of P*xis*, but was otherwise identical to the described constructs.

ELC1760 was used to screen for suppressors of the *orf17*-*16*-caused growth defects. To perform this screen, we wanted two copies of *orf17*-*16* and the genes required for regulation (*immR*, *immA*, and *rapI*) to prevent the acquisition of suppressor mutants with mutations related to *orf17*-*16* expression. Previous constructs were used to generate additional alleles. *lacA*::[(P*xis*- *orf17*-*16*) *mls* (Pxyl-*rapI*) *spc*] was constructed by inserting [(Pxyl-*rapI*) *spc*] into the existing *lacA*::[(P*xis*-*orf17*-*16*) *mls*] allele, selecting for spectinomycin resistance. *amyE*::[(*Pxyl*-*rapI*) *cat* (*PimmR*-(*immR*-*immA*) (*Pxis*-(*orf17*-*16*))] was constructed by inserting *cat*, *PimmR*-(*immR*- *immA*), and *Pxis*-(*orf17*-*16*) into *amyE*::[(*Pxyl*-*rapI*) *spc*] and selecting for chloramphenicol resistance. *yhdGH*::[(P*xis*-*lacZ*) *tetM*] was generated by fusing P*xis*, *lacZ*, and *tetM*, with *yhdG* and *yhdH* homology arms via isothermal assembly, transforming into *B. subtilis*, and selecting for tetracycline resistance.

Δ*yqaR*::*cat* is a deletion-insertion replacing *yqaR* from 18bp upstream of the start of the *yqaR* open reading frame to 48bp upstream of its stop codon with the chloramphenicol resistance cassette from pGEMcat. Fragments were joined via isothermal assembly and used for transformation. A *B. subtilis* strain cured of the *skin* element (leaving behind intact *sigK*) was a generous gift from the Losick lab.

*yhdGH*::[(P*yqaR*-*yqaR*) *kan*] is an insertion of *yqaR* starting 275bp upstream of the start site (to include a putative promoter region) and a kanamycin resistance cassette 19bp downstream of *yhdG* (also called *bcaP*) stop codon. The strong terminator region between *serA* and *aroC* was included upstream of *yqaR* to prevent expression bleed-through from neighboring chromosomal regions. These fragments were joined by isothermal assembly and used for transformation.

Tn*916*-*gfp* was generated as a reporter construct to monitor Tn*916* activation on the single-cell level. It is an unmarked insertion of *gfpmut2* (without a promoter region) 29bp upstream of *orf24*. *Gfpmut2* was fused with up- and downstream-homology into pCAL1422 cut with EcoRI and BamHI. The resulting plasmid, pELC1329, was used to generate ELC1458, which contains a copy of Tn*916*-*gfp* between *yufL* and *yufK* (at GAAAGGGACT TTTT<u>TA</u>TATGAAAAATACTT, where the underlined nucleotides indicate the Tn*916*-chromosome junction). In this context, *gfpmut2* (along with the rest of the DNA processing and T4SS genes) will not be expressed until the element has excised from the chromosome and circularized (Celli and Trieu-Cuot, 1998). To confirm that the growth defects observed upon activation of Tn*916*-*gfp* in ELC1458 were not due to this particular integration site of Tn*916,* this element was subsequently mated into JMA222 (which lacks Tn*916* and ICE*Bs1*) to isolate strains LKM18 and LKM20, which contained Tn*916*-*gfp* at different chromosomal sites: between *nupQ*-*maeN* (TTAGTTTTTT AACT<u>TA</u>AAAA AATATGAAGT) and between *ykuC*-*ykyB* (CAGGTTAAAA ATGC<u>GC</u>TTTT TTTCTTAGAA), respectively. New integration sites were mapped by arbitrary PCR, as previously described (Bean et al., 2021; Brophy et al., 2018; Das et al., 2005).

Tn*916*-*gfp* was transferred via conjugation from *B. subtilis* donor cells into *E. faecalis* (ATCC 19433) recipient cells under standard mating conditions (described below). Briefly, *B. subtilis* Tn*916* donors contained a D-alanine auxotrophy (Δ*alr*::*cat*) to serve as a counter- selection mechanism against donors when selecting for *E. faecalis* Tn*916*-*gfp* transconjugants (Bean et al., 2021; Brophy et al., 2018). To move Tn*916* (Δ*orf17*-*orf16*) into *E. faecalis,* these critical genes were complemented elsewhere in a non-mobile part of the donor chromosome. Tn*916* insertion sites were identified by arbitrary PCR, as above, or by inverse PCR, similar to previously described methods (Lee et al., 2007; Wright and Grossman, 2016). Briefly, chromosomal DNA was digested with either PacI or AseI/NdeI restriction enzymes and then ligated to circularize the fragments. The following primer pairs were used to amplify and sequence the Tn*916*-chromosome junctions: oLM177 (5’- AACGCTTCGT TATGTACCCT CTG) and oLM178 (5’ – ACCACTTCTG ACAGCTAAGA CATG) for PacI digested DNA; oLW443 (5’ – CTCTACGTCG TGAAGTGAGA ATCC) and oLW209 (5’ – TTGACCTTGA TAAAGTGTGA TAAGTCC) for AseI/NdeI digested DNA. Integration sites were mapped to the following sites (with a 30bp genomic context provided, underlined nucleotides indicate the predicted Tn*916*-chromosome junction), of the *E. faecalis* genome (Accession number: ASDA00000000.1). ELC1531 had a copy of Tn*916* between ABC_ACP (TTTTTTACAT GTAT<u>GA</u>TTTT TTTTACAAAA); ELC1529: had two copies: one was between *citG_*SP (AACGGCTGTC GCCT<u>TT</u>TTTT ATGAAATTTT), the second was between HP_CSP (TTTCTTGTTC TTTT<u>TT</u>TTAT AAAAAAAACC); the Δ*orf17*-*orf16* Tn*916* (ELC1696) was between ABH_HP (TCTTTTTTTT GTAA<u>TA</u>AAAA ACAGAAAATT) where ABC = ABC transporter; ACP = acyl carrier protein; SP = surface protein; HP = hypothetical protein; CSP = cold shock protein; ABH = alpha/beta hydrolase (Annotations based on *E. faecalis* KB1 annotations, Accession number: CP015410.1).

### Growth and viability assays

Strains were grown in defined minimal medium with 1% arabinose as a carbon source to early exponential phase. At an OD_600_ of 0.2, the cultures were split into activating or non-activating conditions: 1% xylose was added to stimulate activation of ICE*Bs1*, (ICE*Bs1*-Tn*916*)-H1, or ectopic expression constructs; 2.5 µg/ml tetracycline was added to stimulate activation of Tn*916*. The number of colony forming units (CFUs) was determined immediately prior to activation, and at one or more time points (typically three hours) after activation in induced and non-induced cultures. A “relative viability” was calculated as the number of CFUs present in the induced culture divided by the number of CFUs present in the non-induced culture. OD_600_ was monitored every 30 minutes for four hours post-induction.

### Excision assays

qPCR was used to monitor excision (and therefore activation) of Tn*916*, H1, and ICE*Bs1*, as previously described (Bean et al., 2021; Wright and Grossman, 2016). Briefly, gDNA of ICE host strains was harvested using the Qiagen Dneasy kit with 40 µg/ml lysozyme. The following primers were used to quantify the presence of the empty ICE attachment site, normalized to a nearby chromosomal locus that is present in every cell.

For Tn*916* excision assays (integrated between *yufK* and *yufL*), we used previously described primers (Wright and Grossman, 2016). oLW542 (5’- GCAATGCGAT TAATACAACG ATAC) and oLW543 (5’- TCGAGCATTC CATCATACAT TC) amplified the empty chromosomal attachment site (*att1*). oLW544 (5’- CCTGCTTGGG ATTCTCTTTA TC) and oLW545 (5’- GTCATCTTGC ACACTTCTCT C) amplified a region within the nearby gene *mrpG*.

For H1 and ICE*Bs1* (integrated at *trnS*-*leu2*), oMA198 (5’- GCCTACTAAA CCAGCACAAC) and oMA199 (5’- AAGGTGGTTA AACCCTTGG) amplified the empty chromosomal attachment site (*attB*). oMA200 (5’- GCAAGCGATC ACATAAGGTT C) and oMA201 (5’- AGCGGAAATT GCTGCAAAG) amplified a region within the nearby gene, *yddN*.

qPCR was performed using SsoAdvanced SYBR master mix and the CFX96 Touch Real-Time PCR system (Bio-Rad). Excision frequencies were calculated as the number of copies of the empty chromosomal attachment sites (as indicated by the Cp values measured through qPCR) divided by the number of copies of the nearby chromosomal locus. Standard curves for these qPCRs were generated using *B. subtilis* genomic DNA that contained empty ICE attachment sites and a copy of the nearby gene (*yddN* or *mrpG*). DNA for the standard curves was harvested when these strains were in late stationary phase and had an *oriC*/*terC* ratio of ∼1, indicating that the copy numbers of these targets were in ∼1:1 ratios.

### Suppressor screen

Eighteen independent cultures of ELC1760, which contains two copies of *orf17* and *orf16* (described above, Table 2) were grown in defined minimal medium (with 1% arabinose). In early exponential phase, expression of *orf17* and *orf16* was induced with 1% xylose and cultures were grown overnight (approximately 18 hours) until they became dense. Cultures were back-diluted and this process was repeated 1-2 times to enrich for suppressor mutants. Suppressors were colony-purified, and checked for presence of all antibiotic resistance markers. Additionally, we confirmed these isolates properly activated P*xis*-*lacZ* when streaked on LB plates containing X-gal (5-bromo-4-chloro-3-indolyl-β-D-galactopyranoside) and 1% xylose, indicating that the RapI-driven induction of P*xis* was still working properly (and likely *orf17* and *orf16* were still being expressed).

### Genome resequencing

Each suppressor mutant was grown in a minimal medium containing 1% glucose to an OD_600_ ∼1.5-2 . Cells were harvested, and DNA was extracted using a Qiagen 100 G tips purification kit. Sample preparation, including DNA shearing using a Covaris ultrasonicator, size selection (aiming for ∼500bp), adapter ligation, and paired-end read sequencing (300nt + 300nt) on an Illumina MiSeq were performed by the MIT BioMicro Center. Reads were mapped to the *B. subtilis* JH642 chromosome (Accession number: CP007800) (Smith et al., 2014), as previously described (Deatherage and Barrick, 2014).

### Mating assays

Mating assays were performed essentially as previously described (Auchtung et al., 2005). Briefly, donor strains containing Tn*916* (tetracycline-resistant), (ICE*Bs1*-Tn*916*)- H1 (kanamycin-resistant), or ICE*Bs1*(Δ*rapIphrI*::*kan*) were grown in LB medium to early exponential phase. At an OD_600_ ∼0.2, Tn*916* activation was stimulated with 2.5 µg/ml tetracycline; H1 and ICE*Bs1* activation was stimulated with 1mM IPTG (via the P*spank*(*hy*)-*rapI* allele). After one hour induction, donor strains were mixed in a 1:1 ratio with streptomycin resistant recipient cells (5 total Ods of cells) and filtered. Mating filters were placed on a 1X Spizizen’s salts (Harwood and Cutting, 1990) 1.5% agar plate at 37°C for one hour. Cells were then harvested off the filter and plated on selective media to detect ICE transfer. The number of donor (tetracycline or kanamycin resistant), recipient (streptomycin resistant), and transconjugant (tetracycline/streptomycin resistant or kanamycin/streptomycin resistant) CFUs were enumerated both pre- and post-mating. A conjugation efficiency was calculated as the percentage of transconjugant CFUs formed normalized to the number of donor cells applied to the mating. Conjugation efficiencies were normalized to those of the wild type matings performed in parallel, as described. Typically, conjugation efficiencies were as follows. Tn*916* ∼0.0005%, H1 ∼0.5%, ICE*Bs1* ∼1%.

### Time-lapse microscopy and analysis

*B. subtilis* and *E. faecalis* cells were grown to early-exponential phase in the appropriate medium. When applicable, Tn*916* activation was then stimulated with 2.5 µg/ml tetracycline. After a 1-3 hour induction, cells were transferred to an agarose pad (1.5% UltraPure agarose; Invitrogen) dissolved in growth medium, containing 0.1µg/ml Propidium iodide, DAPI (0.035µg/ml for *B. subtilis*; 0.5µg/ml for *E. faecalis*), and either 2.5 µg/ml tetracycline for Tn*916* activation or 1% xylose for expression of *orf17*-*orf16*- *gfp*. The agarose pad was placed in an incubation chamber, which was made by stacking two Frame-Seal Slide Chambers (Bio-Rad) on a standard microscope slide (VWR). Cells were then grown at 37°C for 2-4 hours while monitoring growth. Fluorescence was generated using a Nikon Intensilight mercury illuminator through appropriate sets of excitation and emission filters (Chroma; filter sets 49000, 49002, and 49008). Time-lapse images were captured on a Nikon Ti- E inverted microscope using a CoolSnap HQ camera (Photometrics). ImageJ software was used for image processing.

## Acknowledgements

We thank Calvin Herman for doing preliminary experiments identifying the growth defects resulting from H1 activation. We thank Laurel Wright for previously constructing the Δ*orf21* deletion used here. We thank Stuart Levine and the BioMicro Center for help with genome resequencing. We thank the Losick lab for sharing a Δ*skin B. subtilis* strain.

Research reported here is based upon work supported, in part, by the National Institute of General Medical Sciences of the National Institutes of Health under award number R35 GM122538 to ADG. Any opinions, findings, and conclusions or recommendations expressed in this report are those of the authors and do not necessarily reflect the views of the National Institutes of Health.

---

**Figure S1.**
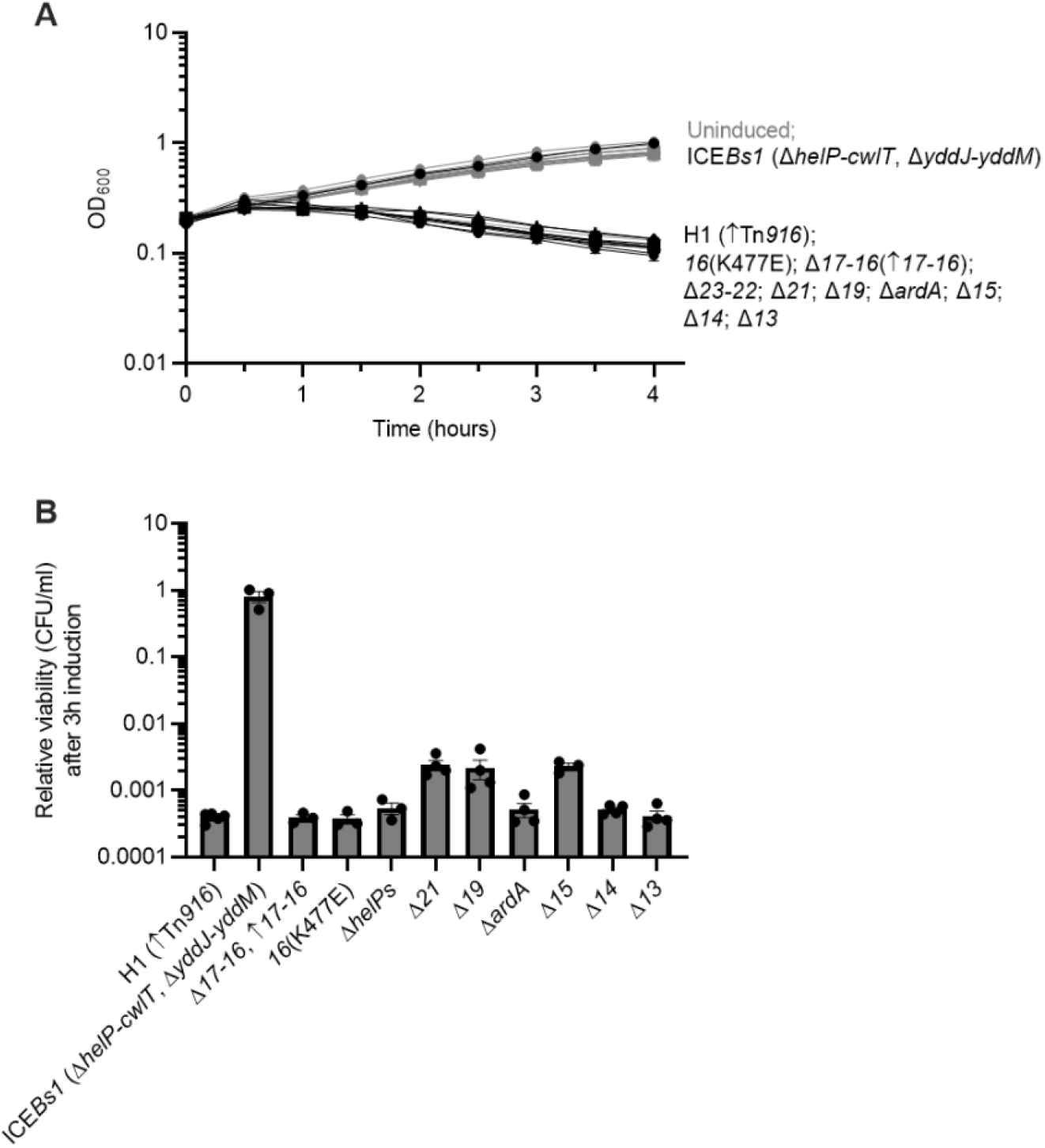
Assessing growth and viability of additional hybrid mutants. Strains containing H1-Δ*attR* (closed squares, ELC1076), ICE*Bs1* (Δ*helP*-*cwlT*, Δ*yddJ*-*yddM*) (circles, ELC1226), or the following H1-Δ*attR* mutants: Δ*orf17*-*orf16* (with *lacA*::P*xis*-*orf17*-*orf16*; plus signs, ELC1938), *orf16*(*K477E*) (asterisks, EL1899), Δ*orf23*-*22* (open upward triangles, ELC1804), Δ*orf21* (downward triangles, ELC1916), Δ*orf19* (hexagons, ELC1915), Δ*ardA* (open squares, ELC1707), Δ*orf15* (open upward triangles ELC1418), Δ*orf14* (open diamonds, ELC1708), and *Δorf13* (closed diamonds, ELC1705) were grown in minimal arabinose medium to early exponential phase. At time = 0 hours, when cultures were at an OD_600_ ∼0.2, cultures were split into inducing (+1% xylose to stimulate *rapI* expression) and non-inducing conditions. **A)** Growth was monitored by OD_600_ for three hours. Black lines indicate growth of the indicated induced cultures; gray lines indicate growth of uninduced cultures. **B)** The relative viability of cultures after three hours of element induction was calculated as the number of colony forming units (CFUs) formed by the induced culture, divided by that from the uninduced culture (a value of “1” indicates there is no change in CFUs with induction). Data presented are averages from three or more independent experiments (individual data points shown in (B)), with error bars depicting standard error of the mean. Error bars could not always be depicted in (A) due to the size of each data point.

## Notes

### Competing Interest Statement

The authors have declared no competing interest.

